# A chromatin-associated pool of Aurora A controls kinetochore-microtubule attachments to ensure chromosome biorientation

**DOI:** 10.1101/2025.10.31.685952

**Authors:** Johnathan L. Meaders, Alyssa A. Rodriguez, Smriti Variyar, SungWoo Park, Alessandro E. Cirulli, Karen Oegema, Kevin D. Corbett, Arshad Desai

## Abstract

Accurate chromosome segregation requires dynamic kinetochore–microtubule attachments that, under the regulation of Aurora family kinases, biorient and align replicated chromosomes. In *C. elegans*, Aurora A acts with the TPX2-related activator TPXL-1 to regulate these attachments and control spindle length. We show that, in addition to prominent spindle pole localization, TPXL-1–AurA has a chromatin-associated pool positioned between the sister kinetochores. Structural modeling and biochemical analysis support TPXL-1 directly recognizing the nucleosome acidic patch via an arginine anchor. Disrupting this interaction selectively removed chromatin-bound TPXL-1–AurA and caused chromosome missegregation, whereas elevation of the chromatin pool disrupted chromosome alignment. These opposing perturbations inversely affected kinetochore recruitment of the microtubule-binding Ska complex. These results support spatially distinct TPXL-1–AurA populations acting sequentially, with the spindle pole pool controlling spindle length by switching kinetochores out of a depolymerization-coupled state, and the chromatin pool controlling attachment stabilization to ensure biorientation prior to anaphase.

## Introduction

Faithful chromosome segregation during cell division is essential for maintaining genomic stability and preventing aneuploidy, a hallmark of cancer and many developmental disorders. During mitosis, replicated sister chromatids are separated by the microtubule-based mitotic spindle, which exerts pulling forces to move the sisters towards opposite poles. Chromosomes assemble kinetochores—large, multiprotein structures that connect centromeric chromatin to spindle microtubules (*1, 2*). Kinetochore-spindle microtubule attachments must be dynamic yet tightly regulated to allow continual error correction until all chromosomes achieve biorientation, the configuration in which each chromatid is connected exclusively to microtubules from one spindle pole and sister chromatids are connected to opposite spindle poles. The coordinated dynamics of kinetochore-microtubule attachments are also central to spindle length regulation and for bioriented chromosomes to align at the spindle midplane to form a metaphase plate. Together, these mechanisms ensure that when cohesion between sister chromatids is dissolved at anaphase onset, the forces transmitted through kinetochore microtubules faithfully segregate one copy of the replicated genome to each daughter cell.

Regulation of kinetochore-microtubule attachments to ensure chromosome biorientation is complex and involves multiple activities, the most prominent of which is phosphorylation by Aurora family kinases—Aurora A (AurA) and Aurora B (AurB) (*3*). Studies in budding yeast, which has a single Aurora kinase, followed by work in metazoans, highlighted a central role for Aurora B (AurB), acting in the context of the chromosomal passenger complex that localizes between the sister kinetochores, in ensuring chromosome biorientation (*4–6*). In metazoans, AurA also plays a key role in kinetochore regulation (*7–14*). AurA exists in multiple pools bound to distinct regulatory factors— including TPX2, CEP192 and BORA in humans—and localizes prominently to spindle poles, mediated by CEP192 and TPX2 (*15*). In addition, AurA has been observed on mitotic chromatin adjacent to kinetochore–microtubule attachments in mammals and in *C. elegans* (*7, 10, 11, 16*), but the function of this chromatin-associated population has remained unclear due to the absence of tools that can selectively perturb it.

Here, we use the *C. elegans* embryo to study the functions of AurA in kinetochore regulation. In *C. elegans*, AurA activated by the TPX2-like cofactor TPXL1 (TPXL1–AurA), regulates kinetochore-microtubule attachments (*8, 12, 17*), whereas AurB plays a minor role (*8*). Depleting TPXL-1 or mutating it to prevent AurA binding traps kinetochores in a persistent depolymerization-coupled state, causing spindle poles to be pulled inward after nuclear envelope breakdown, resulting in a rapid reduction in spindle length (“spindle collapse”); this collapse can be suppressed by preventing outer kinetochore assembly (*8, 12*). These results indicate that TPXL-1–AurA is required for kinetochores to exit from a persistent depolymerization-coupled state upon nuclear envelope breakdown. However, it has remained unclear whether this regulation is mediated by the spindle pole or chromatin-associated pools of AurA, and whether promoting the transition out of the depolymerization-coupled state represents the only function of AurA at kinetochores.

In this study, we define the mechanism by which AurA is recruited to chromatin as well as identify a means to elevate it at this cellular location. By selectively removing or elevating the chromatin-associated pool of AurA, we show that this pool is dispensable for the transition of kinetochores out of the depolymerization-coupled state, suggesting that the spindle pole-associated population is sufficient for this function which is critical for regulation of spindle length. However, we find that the chromatin-associated population is important to ensure biorientation of all chromosomes prior to anaphase onset. We show that the chromatin-associated AurA pool regulates SKA complex recruitment to kinetochores, potentially through phosphorylation of the Ndc80 N-terminal tail, and that its precise level on chromatin is important to biorient and align chromosomes. Together, our findings reveal how spatially distinct pools of TPXL-1–AurA act on kinetochore-microtubule attachments to control spindle length and stabilize bioriented attachments, thereby ensuring accurate chromosome segregation.

## Results

### TPXL-1 directs Aurora A localization to mitotic chromatin

In human cells which have monocentric chromosomes, Aurora A (AurA) localizes not only to spindle poles but also to the inner centromere region between sister kinetochores (*7, 10, 11*). In *C. elegans* embryos, which have holocentric chromosomes, AurA (AIR-1) localizes prominently to spindle poles and has also been reported to localize to chromosomes (*16*). To assess if the mitotic AurA activator TPXL-1 exhibited a similar localization pattern as AurA, we imaged TPXL-1 and AurA fluorescently tagged at their endogenous loci. Both proteins localized robustly to spindle poles (**Fig. 1A**). When the image contrast was adjusted to oversaturate the pole signal, a second pool of TPXL-1 and AurA was observed on mitotic chromosomes (**Fig. 1A; Fig. S1A**). In the *C. elegans* one-cell embryo, the 12 holocentric chromosomes each have two line-shaped kinetochores that extend along the length of the sister chromatids, forming end-coupled interfaces with ∼20 microtubules (*18*). The chromatin-localized TPXL-1 signal overlapped with fluorescent histone H2b and was positioned between sister kinetochores visualized by fluorescently tagging KNL-1, a component of the outer kinetochore scaffold (**Fig. 1B**; (*19*)). These observations identify a TPXL-1–AurA pool on the chromatin between sister kinetochores, analogous to the centromere-associated population of AurA described in human cells (*7, 10, 11*). Thus, the presence of both spindle pole- and chromatin-associated AurA pools appears to be a conserved feature.

**Fig. 1.**
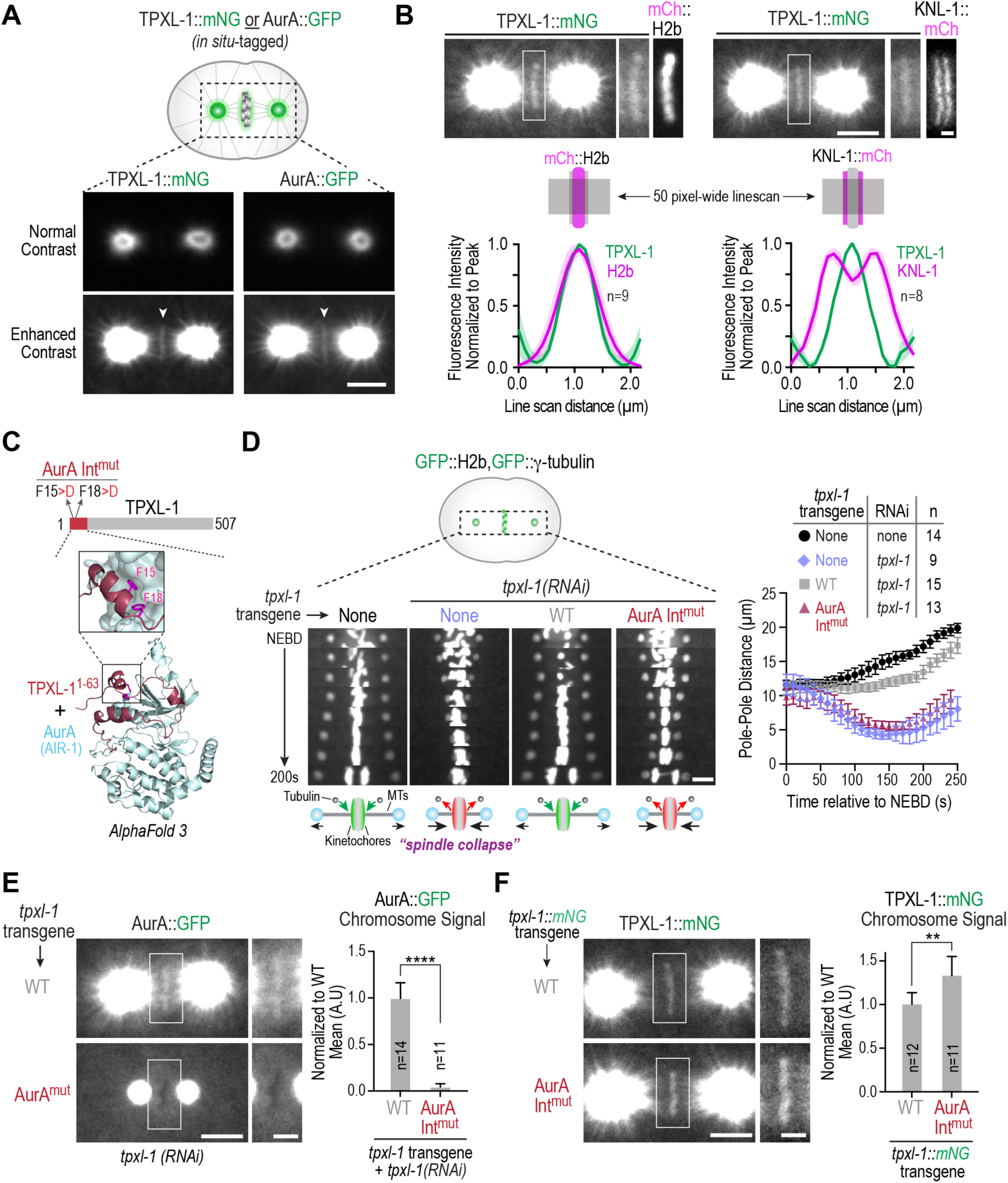
A pool of Aurora A is localized to chromatin via its activator TPXL-1 in one-cell *C. elegans* embryos. **(A)** Images of metaphase embryos expressing indicated *in situ* fluorescent fusions. Enhancing the contrast reveals signal coincident with position of the chromosomes. Scale bar, 5 µm. See also *Fig. S1A*. **(B)** Analysis of *in situ* mNG-tagged TPXL-1 localization relative to mitotic chromatin (marked by mCh::H2b) or kinetochores (marked by KNL-1::mCh). (*top*) Example images; scale bars, 5 µm (*whole spindle*) and 1 µm (*magnified boxed region*). (*bottom*) A 50 pixel-wide linescan was used to measure fluorescence intensities, cytoplasmic background subtracted and signal normalized to the peak prior to averaging across different embryos. Graphs plot the normalized mean signal across the linescan. Shaded region shows the 95% CI. *n* is number of embryos analyzed. **(C)** Alphafold3 model of the N-terminus of TPXL-1 and *C. elegans* Aurora A (AIR-1). The highlighted Phe5 and Phe15 residues of TPXL-1 are targeted for mutation to disrupt its interaction with AurA. See also *Fig. S1B,C*. **(D)** (*left*) Kymographs of the spindle region in strains expressing GFP fusions that mark the chromosomes (GFP::H2b) and the spindle poles (GFP::γ-tubulin). The leftmost column shows a control embryo, while the other columns show TPXL-1 depleted embryos expressing indicated transgenes. Schematics below each column represent the state of kinetochores and of spindle length after NEBD (*MTs: microtubules*). In control (no RNAi and no transgene) or WT TPXL-1 (with endogenous TPXL-1 depletion) conditions, kinetochores are in a polymerization-coupled state (*shaded green*) which enables slow spindle elongation. By contrast, in the absence of TPXL-1 or in the presence of AurAInt^mut^ TPXL-1 (with endogenous TPXL-1 depletion), kinetochores persist in a depolymerization-coupled state (*shaded red*) pulling the spindle poles into the chromosomes (*“spindle collapse”*); preventing outer kinetochore assembly suppresses this spindle collapse phenotype (*8, 12, 17*). Scale bar, 5 µm. (*right*) Graph of spindle pole separation over time for the indicated conditions. *n* is the number of one-cell embryos analyzed. Error bars are SD. **(E) & (F)** Localization analysis of AurA (*E*) and TPXL-1 (*F*) with transgene-encoded WT or the AurA-binding mutant of TPXL-1, following endogenous TPXL-1 depletion. For localization of AurA, the *tpxl-1* transgene insertions were untagged, whereas for TPXL-1 they were mNG-tagged. In both (*E*) and (*F*): (*left*) example images of one-cell embryos for the indicated conditions; the boxed regions are magnified on the right. Scale bars, 5 µm (*whole spindle*) and 2 µm (*magnified boxed region*). (*right*) Graphs of chromosomal signal of AurA (*E*) or TPXL-1 (*F*) for the indicated conditions; measured values were normalized relative to the mean value of the WT condition. *n* is number of embryos analyzed. p<0.0001 (****) and p<0.01 (**) from a Mann-Whitney test. Error bars represent 95% CI.

To determine whether TPXL-1 or AurA mediates localization of the complex to chromosomes, we used a TPXL-1 mutant in which two phenylalanines within its N-terminal AurA-binding domain were mutated to aspartic acid (F15>D;F18>D). In agreement with prior work showing that these residues are required for AurA binding (*12*), Alphafold3 modeling predicted an interface in which the two phenylalanine residues make critical contacts with AurA (**Fig. 1C; Fig. S1B,C**; (*20*)); we thus refer to the AurA interface-disrupting F15>D;F18>D mutant as AurAInt^mut^. Using a single-copy transgene insertion system that allows replacement of endogenous TPXL-1 with transgene-encoded variants (*17*), we confirmed that depleting endogenous TPXL-1 results in a characteristic spindle-collapse phenotype, which likely arises from kinetochore-microtubule attachments persisting in a depolymerization-coupled state after nuclear envelope breakdown (NEBD; **Fig. 1D**). Spindle collapse was rescued by transgenic WT TPXL-1, but not by AurAInt^mut^ TPXL-1, which cannot bind AurA (*12, 17*). In embryos expressing AurAInt^mut^ TPXL-1, AurA no longer localized to chromosomes (**Fig. 1E**), indicating that TPXL-1 recruits AurA to chromosomes. In contrast, AurAInt^mut^ TPXL-1 itself localized to chromosomes (**Fig. 1F**), demonstrating that TPXL-1 binds chromosomes independently of AurA. In the presence of AurAInt^mut^ TPXL-1, AurA was concentrated on the pericentriolar material matrix of the centrosome but failed to extend out from the centrosomes along spindle and astral microtubules (**Fig. 1E**; (*12*)).

Together, these findings define a chromatin-associated pool of TPXL-1–AurA. TPXL-1 associates with chromosomes and recruits AurA to this location, where it would be well-positioned to regulate kinetochore-microtubule attachments during mitosis.

### A short internal region of TPXL-1 is necessary and sufficient for chromatin localization

To explore how TPXL-1–AurA associates with chromatin, we first mapped the region within TPXL-1 required for its chromatin recruitment. Since no recognizable motifs were present following the AurA-binding region at the N-terminus, we employed the transgene-based TPXL-1 replacement system to examine the chromatin localization of truncated variants. This analysis identified an ∼50 amino acid internal region essential for chromatin localization (**Fig. 2A**), which we termed the Chromatin Localization Domain (CLD). Deletion of the CLD from full-length TPXL-1 eliminated chromatin localization but did not affect spindle pole targeting (**Fig. 2B**). TPXL-1 lacking the CLD also failed to recruit AurA to chromatin (**Fig. 2C**), establishing the CLD as being required for the chromatin localization of TPXL-1 and associated AurA.

**Fig. 2.**
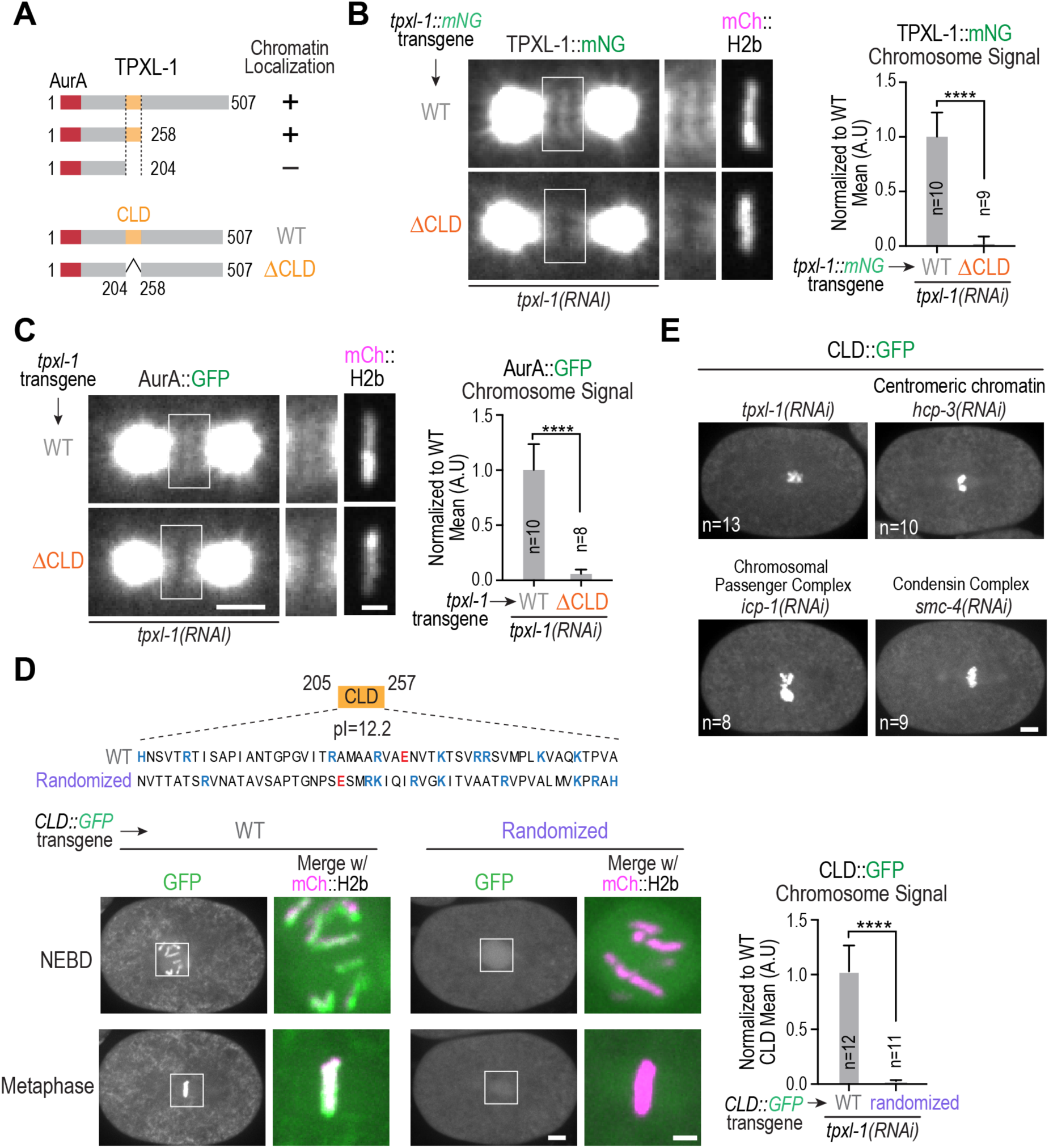
A short internal region of TPXL-1 is necessary and sufficient for chromatin localization. **(A)** (*top*) Summary of TPXL-1 truncation analysis identifying a chromatin localization domain (CLD: aa 205-257). (*bottom*) TPXL-1 transgene-encoded variants employed to analyze the role of the CLD *in vivo*. **(B) & (C)** Localization analysis of transgene-encoded, mNG-tagged WT and ΔCLD TPXL-1 (*B*) or of AurA (*in situ* GFP-tagged) (*C*) in the presence of transgene-encoded, untagged WT and ΔCLD TPXL-1, following endogenous TPXL-1 depletion; the embryos also expressed an mCh::H2b fusion. (*left*) example images of one-cell embryos for the indicated conditions; the boxed regions are magnified on the right; the mCh::H2b region is only shown for the magnified panel. Scale bars, 5 µm (*whole spindle*) and 2 µm (*magnified boxed region*). (*right*) Graphs of chromosomal signal of TPXL-1 (*B*) or AurA (*C*) for the indicated conditions. Error bars are the 95% CI. *n* is number of embryos analyzed. p<0.0001 (****) from a Mann-Whitney test. **(D)** (*top*) Sequence of the 53aa-region CLD highlighting basic (*blue*) and acidic (*red*) residues. A randomized sequence with an identical composition is shown below. (*bottom*) Images of embryos with transgene insertions expressing WT or Randomized CLD::GFP fusions; both NEBD and metaphase stages are shown. The boxed regions are magnified on the right and shown as a merge with mCh::H2b (*magenta*). Scale bars, 5 µm (*whole embryo*) and 2 µm (*magnified boxed region*). (*right*) Graph of chromosomal signal of WT or Randomized CLD; measured values were normalized relative to the mean value of the WT CLD. Error bars are the 95% CI. *n* is the number of embryos analyzed. p<0.0001 (****) from a Mann-Whitney test. See also *Fig. S1D*. **(E)** Images of one-cell embryos expressing the CLD::GFP fusion for the indicated RNAi conditions. *n* is the number of embryos imaged per condition. Scale bar, 5 µm.

To test sufficiency, we expressed the CLD of TPXL-1 as a transgene-encoded fluorescent fusion. The isolated CLD localized robustly and specifically to chromatin, both in the presence and absence of endogenous TPXL-1, confirming that it is sufficient for chromatin association (**Fig. 2D,E; Fig. S1D**). We next investigated candidate mechanisms for how TPXL-1’s CLD directs chromatin localization. Because the CLD is highly basic (pI=12.2), we first tested if its specific sequence or basic character were important. For this purpose, we compared the localization of the wild-type CLD to a sequence-randomized CLD mutant (**Fig. 2D**). While the randomized CLD localized diffusely in the nuclear region, no chromatin localization was observed (**Fig. 2D; Fig. S1D**), highlighting that specific sequence motifs in the TPXL-1 CLD, rather than its overall basic character, are important for chromatin localization.

As TPXL-1 and AurA chromatin signals do not overlap with kinetochores, we did not expect CLD chromatin localization to be impacted by perturbing the centromeric chromatin foundation on which kinetochores assemble. Consistent with this, the TPXL-1 CLD localized robustly to chromatin following depletion of the centromeric histone variant CENP-A (**Fig. 2E**). CLD localization was also unaffected by depletion of the chromosomal passenger complex scaffolding subunit INCENP (ICP-1), or by removal of condensin, via depletion of SMC-4. As cohesin is largely removed from mitotic chromosomes by the prophase pathway in the *C. elegans* embryo (*21, 22*), localization of the TPXL-1 CLD to mitotic chromosomes is also unlikely to depend on cohesin. Thus, TPXL-1’s CLD binds chromatin independently of centromeric chromatin, the chromosomal passenger complex, condensin and, likely, cohesin. These findings suggest that the CLD could directly engage chromatin itself to position a pool of TPXL-1–AurA on chromosomes to regulate kinetochore-microtubule attachments.

### The TPXL-1 chromatin localization domain directly recognizes nucleosomes

The prominent chromatin localization of the TPXL-1 CLD that depends on its specific sequence rather than overall basic character, along with lack of involvement of candidate localization mediators, suggested that the TPXL-1 CLD might directly recognize nucleosomal chromatin. To explore this possibility, we used Alphafold3 to model TPXL-1 with an octameric nucleosome core particle (**Fig. 3A; Fig. S2**; (*20*)). A predicted structure of full-length TPXL-1 with an octameric nucleosome (**Fig. S2A-C**) revealed an interaction between the TPXL-1 CLD and the nucleosome acidic patch, a highly conserved surface on histones H2A and H2B that is a binding site for a range of chromatin-associated factors (*23*). Independent Alphafold3 predictions using the isolated TPXL-1 CLD and either an octameric nucleosome (**Fig. 3A; Fig. S2D-G**) or a H2A-H2B heterodimer identified the same interface with increased confidence (**Fig. S2H-K**). The predicted structure showed TPXL-1 residue R226 docking against several negatively-charged residues on H2A (E61, E64, D90, and E92). This interaction resembles other acidic-patch binders including SIR3 (**Fig. 3B**) and is referred to as the “arginine anchor” (*23*). In addition to the arginine anchor, TPXL-1 residues V223 and M228 are predicted to form hydrophobic interactions with the nucleosome surface. Sequence alignments revealed that all three predicted nucleosome-binding residues (V223, R226, and M228) are conserved in *Caenorhabditis* TPXL-1 homologs (**Fig. 3A**).

**Fig. 3.**
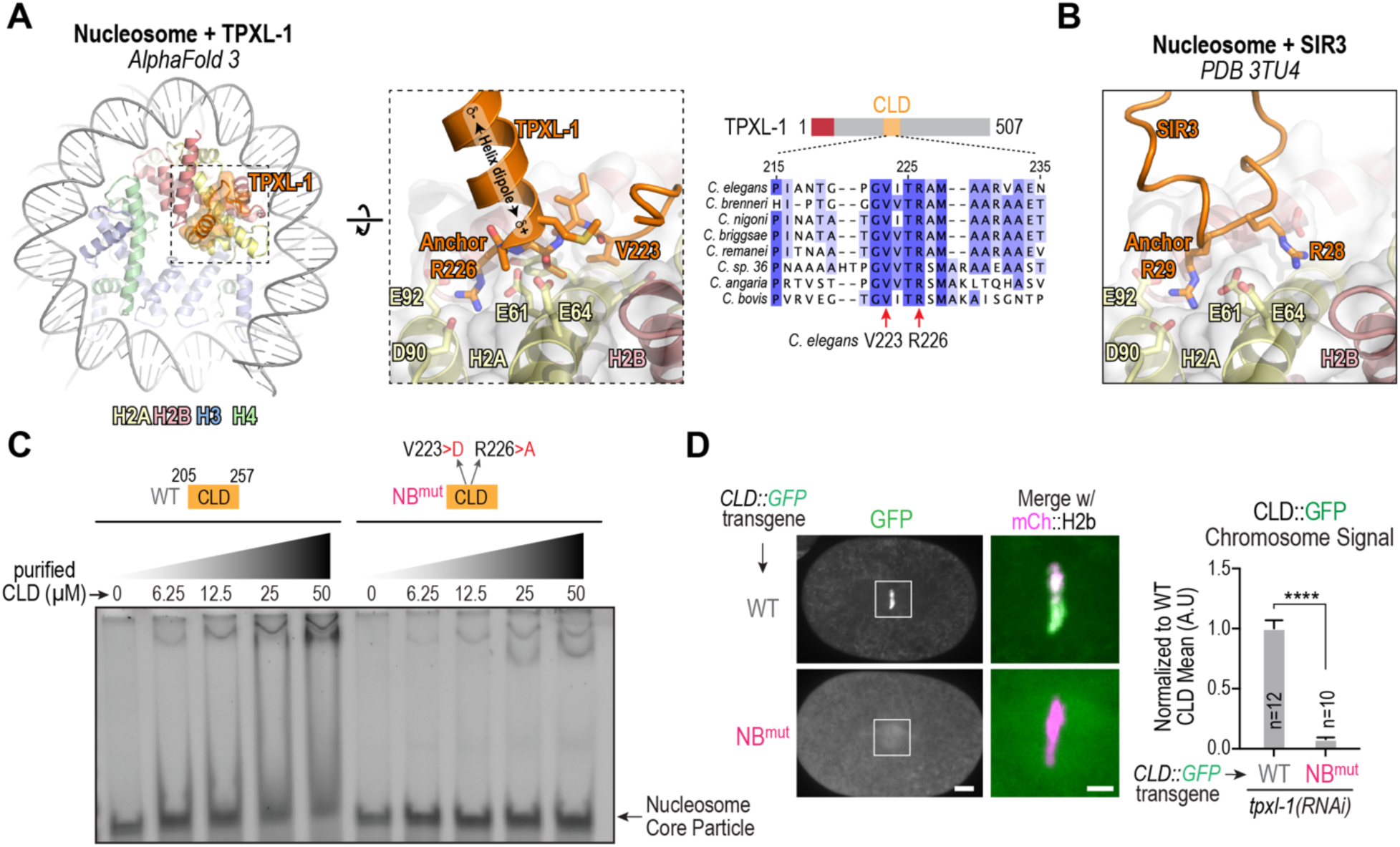
The chromatin localization domain of TPXL-1 directly recognizes nucleosomes. **(A)** (*left*) Alphafold3 model of the central region of the TPXL-1 CLD (aa 217-239) and a nucleosome core particle. *C. elegans* histone sequences and the Widom 601 sequence were employed for the nucleosome model. The model predicts an arginine anchor mechanism for recognition of the nucleosomal acidic patch by the TPXL-1 CLD. The histone residue numbering is based on human histones. See also *Fig. S2*. (*right*) Sequence alignments across related nematode species highlighting conservation of the predicted nucleosome-binding region of the TPXL-1 CLD. The two residues whose mutation are likely to disrupt the predicted interface (V223 and R226) are highlighted. **(B)** A view from the crystal structure (PDB 3TU4) of the interface between the bromo-associated homology (BAH) domain of SIR3 and the nucleosomal acidic patch, which employs an arginine (R29) anchor. **(C)** Nucleosome core particle binding assays with recombinant TPXL-1 CLDs. Both WT and the predicted Nucleosome Binding-mutant (NB^mut^; V223>D; R226>A) were expressed and purified from bacteria. Nucleosome core particles were assembled using *Xenopus* histones and the Widom 601 sequence. Binding was analyzed by an electrophoretic mobility shift assay, employing an increasing concentration gradient of WT or NB^mut^ TPXL-1 CLD. The results shown are representative of two independent experiments. **(D)** Images of embryos with transgene insertions expressing WT or NB^mut^ CLD::GFP. The boxed regions are magnified on the right and shown as a merge with mCh::H2b (*magenta*). Scale bars, 5 µm (*whole embryo*) and 2 µm (*magnified boxed region*). (*right*) Graph of chromosomal signal of WT or NB^mut^ CLD; measured values were normalized relative to the mean value of the WT CLD. Error bars are the 95% CI. *n* is the number of embryos analyzed. p<0.0001 (****) from a Mann-Whitney test.

To test if the TPXL-1 CLD binds nucleosomes, we performed electrophoretic mobility shift assays (EMSAs) using reconstituted *Xenopus* nucleosome core particles and purified recombinant TPXL-1 CLD. Increasing concentrations of WT TPXL-1 CLD caused a concentration-dependent mobility shift of nucleosome core particles (**Fig. 3C**). This shift was abolished by a double mutation of the predicted nucleosome-binding residues V223D and R226A (designated as NB^mut^ for Nucleosome-Binding mutant). To test the functional relevance of this interaction *in vivo*, we generated a strain with an integrated transgene expressing a fluorescent NB^mut^ CLD fusion and analyzed localization. Unlike WT CLD, NB^mut^ CLD did not localize to chromatin (**Fig. 3D**). Together, these results are consistent with the TPXL-1 CLD directly binding to nucleosomes through a conserved arginine-anchor interface, and that this interaction is essential for the TPXL-1 CLD to localize to chromatin *in vivo*.

### The chromatin-associated pool of TPXL-1–AurA is required for accurate chromosome segregation

Having established that the TPXL-1 CLD directly recognizes nucleosomes to recruit TPXL-1– AurA to chromatin, we next asked whether disrupting the nucleosome-binding interface selectively removes TPXL-1–AurA from chromatin *in vivo* and whether this affects spindle assembly and chromosome segregation. Using the TPXL-1 replacement system, we expressed GFP-tagged and untagged versions of WT and NB^mut^ TPXL-1. NB^mut^ TPXL-1 failed to localize to chromatin but retained robust spindle pole localization (**Fig. 4A**; **Fig. S3A**). Correspondingly, AurA was selectively lost from chromatin in NB^mut^ TPXL-1 embryos, while its spindle pole localization appeared unchanged (**Fig. 4A**). Quantitative analysis confirmed this selective depletion (**Fig. 4A**), indicating that the 2-residue mutation that disrupts nucleosome binding by the TPXL-1 CLD removes the chromatin-associated pool of TPXL-1–AurA without affecting its spindle pole localization *in vivo*.

**Fig. 4.**
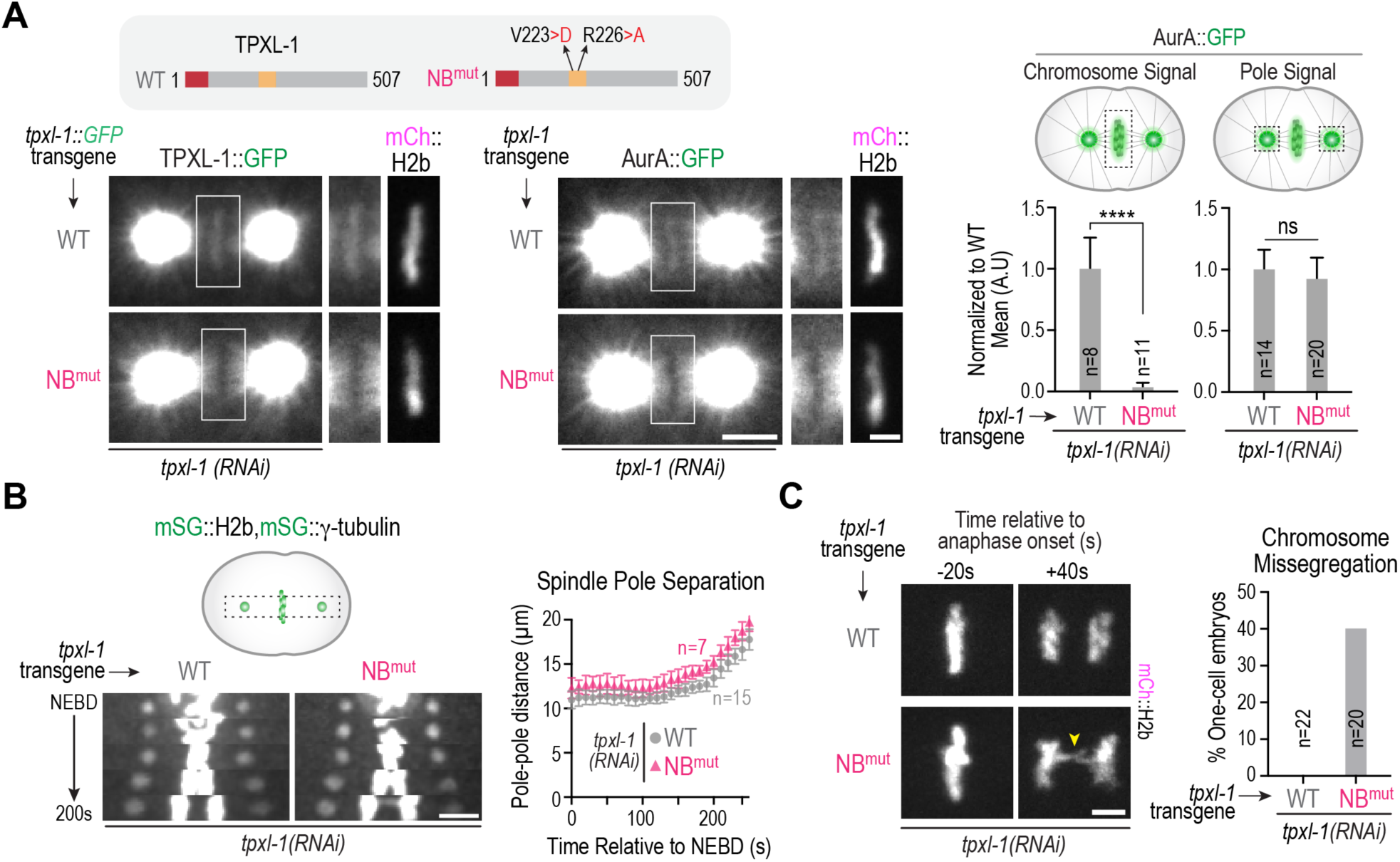
Functional consequences of selectively removing the chromatin pool of TPXL-1–AurA. **(A)** (*top gray box*) Schematics of WT and NB^mut^ TPXL-1 expressed from transgene insertions, either with or without a GFP tag. (*bottom left*) Localization of transgene-encoded TPXL-1::GFP, either WT or NB^mut^, following endogenous TPXL-1 depletion. (*bottom right*) Localization of AurA::GFP in the presence of untagged, transgene-encoded WT or NB^mut^ TPXL-1, following endogenous TPXL-1 depletion. The boxed regions are magnified on the right; the mCh::H2b region is only shown for the magnified panel. Scale bars, 5 µm (*whole spindle*) and 2 µm (*magnified boxed region*). (*right*) Graphs plotting AurA::GFP signal on chromosomes (*left graph*) or on spindle poles (*right graph*) for the indicated conditions. Measured values were normalized relative to the WT mean value. Error bars are the 95% CI. *n* is the number of embryos analyzed per condition. p<0.0001 (****) and not significant (ns) from Mann-Whitney tests. See also *Fig. S3A*. **(B)** (*left*) Kymograph of spindle region of embryos expressing monomeric StayGold (mSG) fusions to label chromosomes (H2b) and spindle poles (γ-tubulin). The indicated *tpxl-1* transgene insertions were present and endogenous TPXL-1 was depleted. Scale bar, 5 µm. (*right*) Graph of spindle pole separation over time for the indicated conditions. *n* is the number of one-cell embryos analyzed. Error bars are SD. See also *Fig. S3B*. **(C)** (*left*) Images from time lapse movies of chromosomes in metaphase and 40s after anaphase onset, for the indicated conditions. Scale bar, 2.5 µm. (*right*) Graph plotting the percentage of one-cell embryos with chromatin bridges in anaphase for the indicated conditions. Data shown pools analysis of WT and NB^mut^ TPXL-1 in mCh::H2b and mSG::H2b expressing embryos. See also *Fig. S3C*.

To assess the functional significance of the chromatin pool of TPXL-1–AurA, we introduced fluorescent markers to visualize chromosomes and spindle poles and analyzed phenotypes following endogenous TPXL-1 depletion in embryos expressing transgenic WT or NB^mut^ TPXL-1. In contrast to embryos depleted of TPXL-1 with no replacement or expressing the AurA-binding mutant, which exhibited spindle collapse (**Fig. 1D**), transgenic NB^mut^ and ΔCLD TPXL-1 supported spindle elongation equivalently to WT TPXL-1 (**Fig. 4B**; **Fig. S3B**). Thus, the chromatin-associated TPXL-1– AurA pool is not required for the transition of kinetochores out of the depolymerization-coupled state, suggesting that the spindle pole-localized population is sufficient for this function. Notably, however, elevated rates of chromosome missegregation, as measured by the frequency of anaphase chromatin bridges, were observed in both NB^mut^ and ΔCLD TPXL-1 embryos (**Fig. 4C**; **Fig. S3C**). These bridges are characteristic of defective biorientation of the holocentric *C. elegans* chromosomes, suggesting that the chromatin pool of TPXL-1–AurA ensures chromosome biorientation on the spindle, potentially by regulating kinetochore-microtubule attachments. Together, these results suggest that distinct populations of TPXL-1–AurA carry out separable functions—the population concentrated at and emanating out from spindle poles is sufficient to promote kinetochores to exit from a depolymerization-coupled state and to regulate spindle length, and the population on chromatin ensures bioriented kinetochore-microtubule attachments.

### A short C-terminal deletion of TPXL-1 elevates the chromatin-associated pool of TPXL-1– AurA

Imaging of *in situ*-tagged TPXL-1 and AurA revealed that while a small fraction of TPXL-1– AurA localizes to mitotic chromatin, the majority is concentrated at the spindle poles. In contrast, the isolated TPXL-1 CLD localized robustly to mitotic chromatin, indicating that the limited chromatin pool of TPXL-1–AurA is not due to a shortage of nucleosomal binding sites. This observation suggested that additional regions of TPXL-1—or its association with other structures such as spindle poles or microtubules—may restrict CLD-mediated accumulation on chromatin. We therefore asked whether the precise amount of chromatin-bound TPXL-1–AurA is important for ensuring accurate chromosome segregation. Insight into this question came from the truncation analysis of TPXL-1, which revealed that a short (50 aa) C-terminal deletion of TPXL-1 (ΔC50) significantly elevated (by ∼3-4 fold) the amount of TPXL-1 (**Fig. 5A**) and AurA (**Fig. 5B**) on chromosomes. Although the mechanistic basis for this enhanced chromatin association remains to be clarified, the ΔC50 mutant provided a useful tool to assess the consequences of elevating the chromatin-associated AurA pool, complementing the nucleosome-binding mutant that removes this pool.

**Fig. 5.**
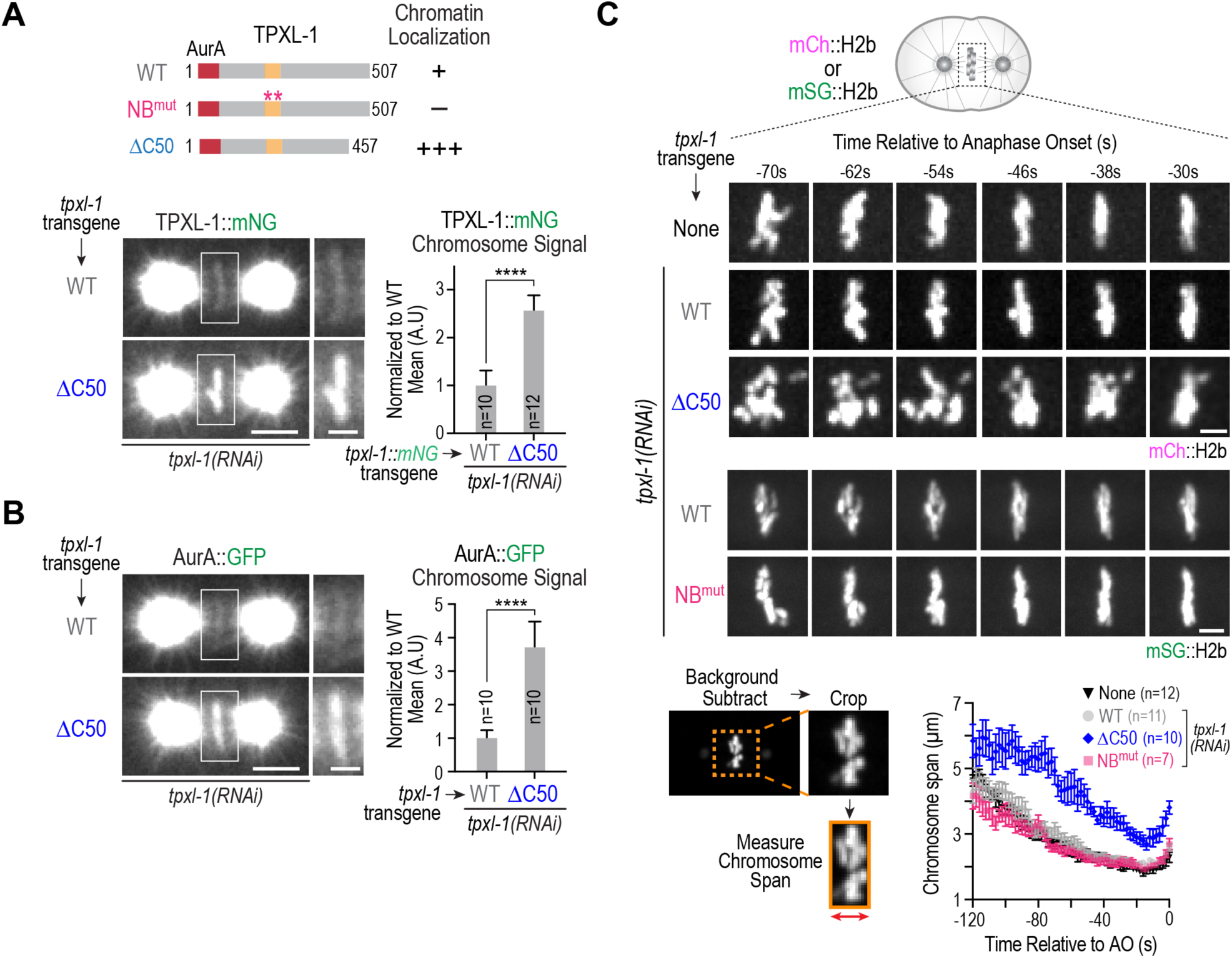
A short C-terminal deletion of TPXL-1 elevates chromatin localization and impairs chromosome alignment. **(A)** (*top*) Schematic summary of chromatin localization analysis of indicated TPXL-1 variants. (*bottom*) Example images of one-cell embryos expressing WT or ΔC50 TPXL-1::mNG following endogenous TPXL-1 depletion; the boxed regions are magnified on the right. (*right*) Graph of chromosomal signal of WT and ΔC50 TPXL-1; measured values were normalized relative to the mean value of the WT condition. See also *Fig. S3D*. **(B)** Analysis of AurA localization for the indicated conditions; the boxed regions are magnified on the right. (*right*) Graph of chromosomal AurA signal for the indicated conditions; measured values were normalized relative to the mean value of the WT condition. Scale bars in panels (*A*) and (*B*): 5 µm (*whole spindle*) and 2 µm (*magnified boxed region*). Error bars in graphs in (*A*) & (*B*) are the 95% CI; *n* is number of embryos analyzed; p<0.0001 (****) from a Mann-Whitney test. **(C)** (*top*) Timelapse imaging of chromosome dynamics for the indicated conditions in embryos expressing either mCh::H2b (*top 3 conditions*) or mSG::H2b (*bottom 2 conditions*). Image panels are from movies that were time-aligned relative to anaphase onset. Scale bars, 2.5 µm. (*bottom left*) Schematic of method used to measure chromosome span on the spindle over time; (*bottom right*) Graph plotting chromosome span over time for the indicated conditions. Error bars are the SEM. The WT condition pools measurements from timelapse sequences obtained from mCh::H2b and mSG::H2b expressing embryos; separated WT traces for each strain background are shown in *Fig. S3E*.

Embryos expressing ΔC50 TPXL-1 did not exhibit spindle collapse after NEBD (**Fig. S3D**). However, analysis of chromosome behavior revealed a distinct defect compared to NB^mut^ TPXL-1 (**Fig. 5C**). In embryos expressing WT or NB^mut^ TPXL-1, chromosomes coalesced onto the metaphase plate and largely ceased their motion along the spindle axis over the interval between 120 and 60 seconds prior to anaphase onset (**Fig. 5C**, *−120 to −60s*), followed by a small amount of additional tightening of the plate during the interval between 60 and 20 seconds prior to anaphase onset (**Fig. 5C**, *−60 to −20s*). In contrast, ΔC50 TPXL-1 embryos failed to coalesce their chromosomes onto the metaphase plate during the −120 to −60s interval. Chromosomes instead remained highly dynamic along the spindle axis, resulting in a wider, disorganized chromosome configuration that persisted through late prometaphase and metaphase (**Fig. 5C**). Quantification of chromosome span (the width of the chromosome mass along the spindle axis; (*8*)) over time underscored this effect (**Fig. 5C; Fig. S3E**). In control, WT TPXL-1 and NB^mut^ TPXL-1 embryos, chromosome span decreased from ∼4.5 to ∼2.5 µm during the - 120 to −60s interval as chromosomes aligned at the metaphase plate and ceased their movements along the spindle axis (**Fig. 5C**). In contrast, the chromosomes in TPXL-1 ΔC50 embryos exhibited a span of 4.5 µm at the end of the −120 to −60s interval, reflecting persistent chromosome movement and incomplete congression; while a reduction in chromosome span was observed over the interval between −60 and −20s, the chromosomes never fully congressed to form a tight metaphase plate (**Fig. 5C**).

Together, these findings indicate that the level of chromatin-associated TPXL-1–AurA must be precisely balanced: a low level of chromatin-associated TPXL-1–AurA is required to facilitate biorientation, likely through control of maturation of kinetochore-microtubule attachments, while excess chromatin-associated TPXL-1–AurA interferes with attachment stabilization, preventing chromosome alignment before anaphase onset.

### Elevating TPXL-1–AurA on chromatin phenocopies loss of SKA complex function

The delayed alignment and persistent chromosome motion observed in ΔC50 TPXL-1 embryos—which have elevated levels of TPXL-1–AurA on chromatin— resembled the phenotype caused by inhibiting the microtubule-binding SKA complex (*8*). In *C. elegans*, the SKA complex is recruited in an NDC-80 complex-dependent manner and reduces the dynamicity of kinetochore-microtubule attachments to align chromosomes prior to anaphase onset (*8*). To compare the ΔC50 TPXL-1 and SKA complex loss phenotypes directly, we quantified chromosome span over time. Chromosomes in ΔC50 TPXL-1 and *ska-3(RNAi)* embryos exhibited very similar dynamics, confirming the similarity of the effects of the two perturbations (**Fig. 6A**; **Fig. S4A**). Consistent with prior work (*8*), imaging of *in situ* tagged SKA-1 revealed that it normally becomes prominent at kinetochores during a short window before anaphase, coinciding with dampened chromosome motion and tight metaphase plate formation (**Fig. 6B**). We next examined the effect of ΔC50 TPXL-1 on SKA-1 recruitment. As shown previously (*24*), TPXL-1 depletion or mutation of its AurA binding site caused hyper-recruitment of SKA to chromosomes (**Fig. 6C**). Expression of WT TPXL-1 restored SKA-1 recruitment to normal levels, whereas ΔC50 TPXL-1 significantly reduced SKA localization to kinetochores (**Fig. 6C**), which likely explains the persistent chromosome movement and incomplete alignment observed when chromosomal AurA is elevated. To complement this analysis, we also analyzed the effect of removing the chromatin-associated TPXL-1–AurA pool using NB^mut^ TPXL-1. In this case, a modest but significant increase in SKA complex recruitment was observed (**Fig. 6C**), suggesting that loss of the chromatin associated TPXL-1–AurA pool enhances SKA loading, potentially leading to premature stabilization of erroneous attachments and the observed chromosome missegregation. The lower extent of SKA over-recruitment seen with the NB^mut^ TPXL-1 compared with TPXL-1 depletion or loss of TPXL-1 AurA binding (**Fig. 6C**) suggests that TPXL-1–AurA emanating out from the spindle poles also restrains SKA recruitment to kinetochores. Together, these findings indicate that the chromatin-associated pool provides an additional layer of SKA recruitment control that is important to biorient all chromosomes.

**Fig. 6.**
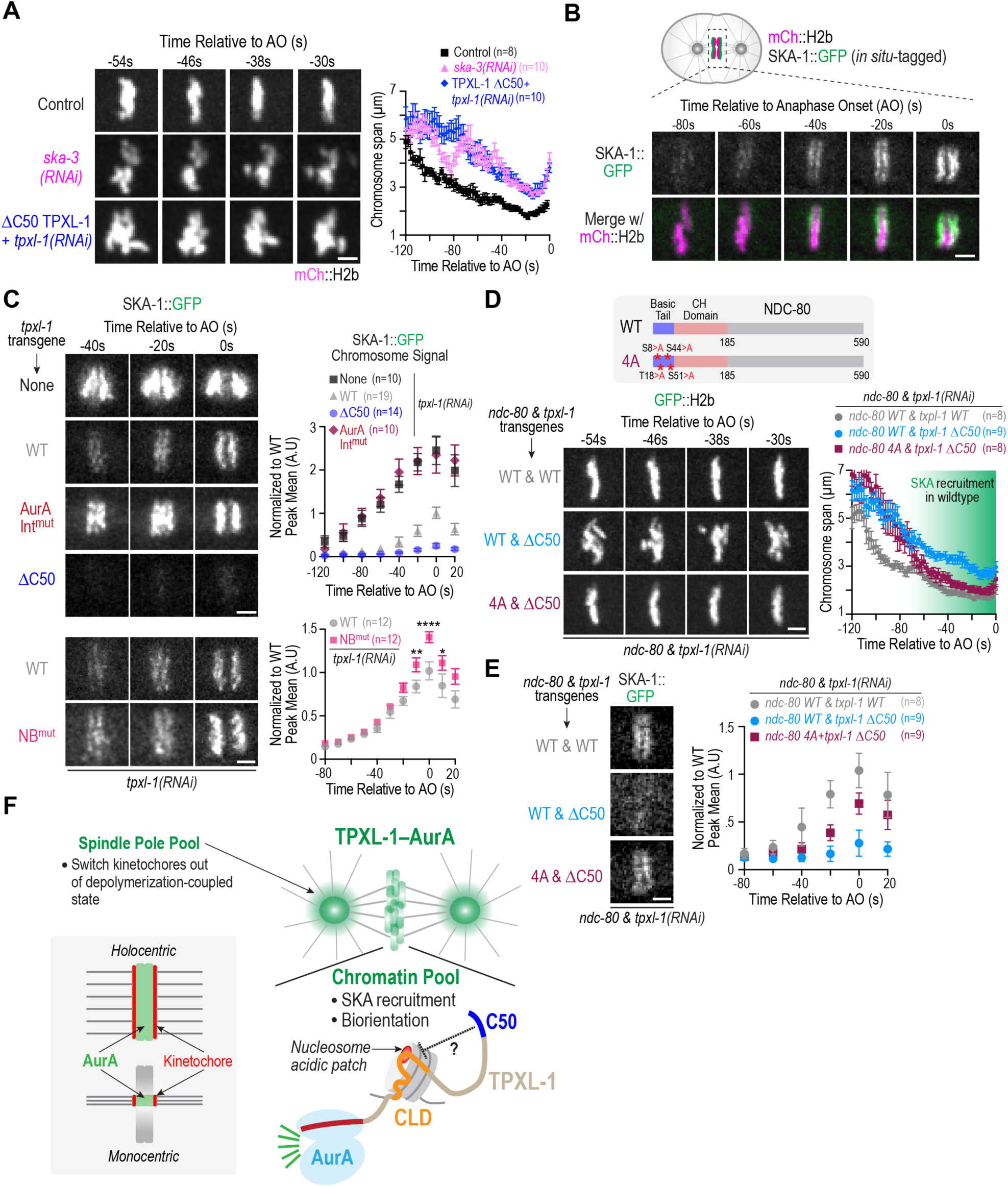
Chromatin-localized AurA regulates the Ndc80-Ska module at kinetochores. **(A)** Example image sequences and quantification of chromosome span over time, performed as in Fig. 5C, for the indicated conditions. The ΔC50 TPXL-1 data plotted in the graph is the same as in Fig. 5C. The Control curve is of unperturbed embryos expressing mCh::H2b. Scale bar, 2.5 µm. Error bars are the SEM. *n* is the number of embryos analyzed. See also *Fig. S4A*. **(B)** Image panels from a timelapse movie highlighting progressive loading of the SKA complex onto kinetochores during mitosis. *In situ* GFP-tagged SKA-1 was imaged along with mCh::H2b as a chromosomal marker. Scale bar, 2.5 µm. **(C)** Analysis of SKA complex localization for the indicated TPXL-1 variants following endogenous TPXL-1 depletion. Image panels from timelapse movies, aligned relative to anaphase onset, are shown on the left. Scale bars, 2.5 µm. Quantification of chromosomal SKA-1::GFP signal over time is shown on the right. Error bars are the 95% CI; *n* is the number of embryos analyzed. In the bottom graph, p-values are from Mann-Whitney tests; **** (p<0.0001), ** (p=0.0068), * (p=0.045). **(D)** Analysis of the effect of the NDC-80 4A N-tail phosphosite mutation (*schematic on top*) on chromosome dynamics in WT versus ΔC50 TPXL-1 variants. Image panels show chromosome behavior over time for the indicated conditions; graph on the right plots the chromosome span over time, measured as in Fig. 5C. *n* is number of embryos analyzed; error bars are the SEM. Scale bar, 2.5 µm. See also *Fig. S4B*. **(E)** Analysis of SKA complex localization for the indicated conditions. Image panel shown is just prior to anaphase. Scale bar, 2.5 µm. Graph on the right plots chromosomal SKA-1::GFP signal over time. Error bars are the 95% CI; *n* is number of embryos analyzed. **(F)** Schematic summary of key findings. TPXL-1–AurA is present in chromatin-associated and spindle pole-associated pools. Chromatin association is likely mediated by direct recognition of the nucleosome acidic patch by an arginine anchor in the chromatin localization domain (CLD) that is in the middle of TPXL-1; the N-terminal AurA binding region of TPXL-1 binds and activates AurA. The C-terminal 50 amino acids (C50) of TPXL-1 restrain its chromatin localization by an unknown mechanism. Selective manipulation of the chromatin pool of TPXL-1–AurA impacts SKA complex recruitment, and this pool is required for ensuring biorientation of all chromosomes. The chromatin pool is, however, dispensable for switching kinetochores out of a depolymerization-coupled state, indicating that this function must be provided by the spindle pole pool. Inset panel highlights presence of AurA on chromatin adjacent to kinetochore-microtubule attachments in holocentric (*C. elegans*) and monocentric (vertebrate) species.

The observation that chromatin-associated TPXL-1–AurA negatively regulates SKA complex recruitment raised the question of which AurA target sites mediate this effect. Prior work has shown that mutating four Aurora kinase phosphorylation sites in the basic N-terminal tail of Ndc80 (Ndc80 4A mutant) increases SKA complex recruitment (*8*). To test whether phosphorylation at these sites is required for the chromatin-associated TPXL-1–AurA pool to exert its effect on chromosome dynamics, we generated strains combining the transgenes expressing WT or ΔC50 TPXL-1 with transgenes expressing WT or non-phosphorylatable 4A NDC-80 and analyzed chromosome dynamics following endogenous TPXL-1 and NDC-80 depletion. When NDC-80 4A was present, the persistent chromosome movement and incomplete congression as well as the reduced SKA complex recruitment caused by ΔC50 TPXL-1 were significantly rescued (**Fig. 6D,E; Fig. S4B**). These results are consistent with chromatin-associated TPXL-1–AurA regulating SKA complex recruitment and chromosome dynamics through phosphorylation of the NDC-80 N-terminal tail.

Overall, the above data indicate that the chromatin pool of TPXL-1–AurA controls SKA complex recruitment to stabilize kinetochore-microtubule attachments, with a precise level of TPXL-1–AurA activity at this location being critical to ensure biorientation of all chromosomes prior to anaphase onset.

## Discussion

Mitotic kinases of the Aurora, Plk (Polo-like kinase) and Cdk (cyclin-dependent kinase) families act on hundreds of substrates in distinct cellular locations to ensure that the replicated genome is accurately distributed to daughter cells. A major challenge with investigating mitotic kinase functions is their breadth of action, which leads to complex phenotypes following kinase activity inhibition. Here, we focus on Aurora A, a multi-functional kinase present in different regulator-bound pools concentrated at diverse cellular locations. Our analysis focused on the AurA populations associated with TPXL-1, an ortholog of TPX2, that concentrates at spindle poles and on chromatin and is a major regulator of spindle assembly and chromosome segregation. Our findings reveal that spatially distinct pools of TPXL-1–AurA perform complementary functions in regulating kinetochore–microtubule attachments that in turn impact spindle length and chromosome biorientation. The spindle pole– associated TPXL-1–AurA population promotes the transition of kinetochores out of a depolymerization-coupled state following nuclear envelope breakdown, enabling the proper regulation of spindle length. The chromatin-associated pool functions as a brake on SKA complex recruitment, enabling resolution of attachment errors and full biorientation (**Fig. 6F**). While the pool at spindle poles also restrains SKA recruitment, we suggest that the chromatin pool provides a second layer of control that is critical to ensure biorientation as chromosomes move away from the poles on their path to alignment on the metaphase plate.

Our ability to address the functions of distinct TPXL-1–AurA pools was facilitated by the elucidation of the mechanism by which it is recruited to chromatin. We found that TPXL-1 directly recognizes nucleosomes, likely employing an arginine anchor mechanism to engage the nucleosome acidic patch (**Fig. 6F**). This mechanism of nucleosome recognition is widespread and has been characterized in many diverse factors that engage with chromatin (*23*). Engineering mutations predicted on the basis of structural modeling to disrupt nucleosome recognition by TPXL-1 abrogated nucleosome binding *in vitro* and chromatin localization *in vivo*. Removal of the chromatin pool of TPXL-1–AurA did not cause spindle collapse, indicating that the spindle pole pool is sufficient to switch kinetochores out of a depolymerization-coupled state and allow assembly of a normal length spindle. Instead, loss of the chromatin pool resulted in anaphase chromosome bridges. In the *C. elegans* embryo, which have holocentric chromosomes, anaphase bridges are characteristic of biorientation defects where the kinetochore of a single chromatid connects to microtubules from both spindle poles.

Notably, the loss of the chromatin pool elevated SKA complex recruitment suggesting that incorrect kinetochore-microtubule attachments are prematurely stabilized, leading to the anaphase defects. Conversely, analysis of a mutant that elevated the TPXL-1–AurA chromatin pool prevented SKA complex recruitment, resulting in a kinetochore-attachment stabilization defect and hyperdynamic chromosome movement on the spindle. In the *C. elegans* embryo, the cessation of chromosome movement coincident with alignment to the equator is dependent on SKA recruitment by the NDC-80 complex (*8*). In this system, there is no equivalent of the Astrin-SKAP complex that lubricates the bioriented kinetochore-microtubule interface in human cells (*25*); consequently, chromosome behavior in the *C. elegans* embryo resembles what is observed in human cells lacking Astrin-SKAP.

The effects of loss of SKA function are observed during the interval between −120s and −60s prior to anaphase onset, when levels of SKA on chromosomes are relatively low, suggesting that biorientation and alignment are mediated by low-level SKA recruitment. A major wave of SKA recruitment occurs after chromosome alignment during the 60s interval immediately prior to anaphase onset; this likely represents a second step in attachment stabilization that prepares kinetochores to withstand pulling forces after cohesion is lost in anaphase. A question raised by these observations is how a small, constant level of TPXL-1–AurA associated with chromatin precisely regulates SKA recruitment so that stabilization is coupled to the success of chromosomes in achieving a bioriented state. As SKA recruitment is regulated by NDC-80 phosphorylation, we speculate that the dynamic recruitment or regulation of phosphatase activities may play a key role in sensing biorientation. One possibility is that phosphorylation of targets like the N-terminal tail of NDC-80 by chromatin-bound TPXL-1–AurA holds kinetochore-microtubule attachments in a dynamic state until the threshold set by this phosphorylation is overcome by phosphatases recruited in response to biorientation and alignment. One of the major phosphatases at kinetochores is PP1, which is recruited to its docking site on the KNL-1 kinetochore scaffold during this interval (*26*). However, loss of KNL-1 docked PP1 does not cause the persistent hyperdynamic chromosome movement phenotype (*19*), suggesting that regulation of TPXL-1–AurA activity or of other phosphatases may also contribute to coupling kinetochore-microtubule attachment stabilization to biorientation.

TPXL-1 is divergent from TPX2, its vertebrate ortholog, and chromatin association of TPX2 has, to our knowledge, not been reported. Alphafold3 modeling also does not predict interaction of human TPX2 with nucleosomes. However, multiple studies in human cultured cells have reported AurA targeting to the inner centromere, the chromatin region between the sister kinetochores on monocentric chromosomes, and association of AurA with chromosomal passenger complex subunits has been suggested to mediate this localization (*7, 10, 11*). In additional functional studies have implicated Aurora A in spindle length control and kinetochore-microtubule attachment regulation (*9–11, 13, 14, 27*). Thus, while the precise mechanism of chromatin targeting may differ, in both holocentric *C. elegans* and monocentric human cells, AurA activity is concentrated on the chromatin adjacent to kinetochore-microtubule attachments (**Fig. 6F**). We speculate based on our results that, across this large evolutionary distance, AurA is present at this site because the balance between local AurA and opposing phosphatase activities controls the maturation of kinetochore-microtubule attachments via phosphoregulated SKA complex recruitment, which in turn stabilizes the bioriented configuration prior to anaphase onset.

## Materials and Methods

### C. elegans Strains

*C. elegans* strains used in the study are listed in Table S1. All strains were maintained on NGM plates seeded with OP50 *Escherichia coli* to produce a feeding lawn. Hermaphrodites were periodically passed to fresh plates to prevent starvation and stored at 20°C.

All RNAi-resistant transgenes were generated by single-copy insertions at specific chromosomal loci using the MosSCI method (*28*). Transgene insertions expressing TPXL-1 WT and AurA binding mutants, tagged with mNG or untagged, were previously described (*17*). All other TPXL-1 mutants used in this study were generated by injecting both gonad arms of EG6429 young adults with pCFJ350 integration plasmids containing the TPXL-1 mutations indicated in each figure. Gene blocks containing the nucleosome binding mutants (V233>D;R226>A) were purchased from IDT and Gibson cloned into pCFJ350. TPXL-1::GFP fusions contain GFP with introns enriched for periodic An/Tn clusters to prevent transgene silencing (referred to as GFP[PATC-enriched]) (*29*). NDC-80 RNAi-resistant strains were previously described (*8*).

Integration plasmids (pCFJ350) were mixed with co-injection markers pCFJ190, pCFJ104, pGH8, and pMA122 to screen against extrachromosomal arrays, along with a transposase (pCFJ601). Injected worms were singled onto NGM plates and grown for 7-10 days before heat shock was performed for 3 hrs at 34°C. Surviving progeny were screened for the absence of array markers (mCherry), and singled onto fresh plates. Successful integration was assessed by PCR, and integrated cassettes were sequence-validated using Oxford Nanopore longread sequencing.

Spindle poles and chromosomes were tagged using monomeric Staygold (mSG; (*30, 31*)) fused to the C-terminus of *tbg-1* and to the N-terminus of *his-11* in the same operon, separated by an operon linker. The operon cassette was then integrated on chromosome I using the MoSCI method.

In situ GFP-tagged *ska-1* was previously described (*8*). Endogenous *tpxl-1* was tagged at the C-terminus using CRISPR/Cas9 (*32*). A Cas9-RNP mix, containing the guide sequence (5’-CCCGGGCACTGCTTCGA-3’), repair template, Cas9 (purchased from Berkley MacroLab), and selection markers were injected into young N2 adult hermaphrodites. After 3-5 days, successfully edited strains were validated using genotyping PCR and Sanger sequencing of the edited genomic region.

### RNAi

Double stranded RNA (dsRNA) was produced by amplifying DNA templates from *C. elegans* cDNA or genomic DNA using primers containing the T7 and T3 promoter sequences listed in Table S2 for PCR. Templates were purified using a QIAquick PCR Purification Kit (Qiagen). Single stranded RNA was generated from DNA templates using MEGAscript T3 and T7 Transcription Kits (Invitrogen) and purified using a MEGAclear Transcription Clean-up Kit (Invitrogen). Single stranded RNA was subsequently annealed at 68°C for 10 minutes followed by 37°C for 30 minutes to produce final dsRNA products. L4 hermaphrodites were injected with final dsRNA products. For tpxl-1*(RNAi)*, injected animals were incubated at 16°C for 24 hrs, followed by a shift to 20°C for 24 hrs before imaging. L4’s injected with *hcp-3*, *icp-1*, and *smc-4*(*RNAi)* (**Fig. 2E**) and *ska-3(RNAi)* (**Fig. 6A**) were incubated at 20°C for 38-46 hrs before dissection and imaging. Co-depletion of endogenous *tpxl-1* and *ndc-80* (**Fig. 6D** and **5E**) was performed by mixing each dsRNA at 1:1 ratios and incubating L4 hermaphrodites at 20°C for 48 hrs before imaging.

### Fluorescence imaging of C. elegans embryos

Embryos were dissected from adult hermaphrodites into M9 buffer and mounted on a 2% agarose pad using a mouth pipette. Embryos were then covered with a 22X22 mm coverslip (No. 1.5), and all imaging was performed in a room cooled to 20°C. Spindle pole separation, chromatin localization, and kinetochore recruitment assays were imaged on a Nikon Ti2 confocal system with a Yokogawa CSU-X1 confocal scanner and an Andor iXon electron multiplication back-thinned charge-coupled device (EMCCD) camera (**Fig. 1D** and **S3B,D**) or a Kinetix 22 active-pixel sensor (CMOS) (**Fig. 4B**) using a 60x 1.42 NA Plan Apochromat Lens (Nikon), using NIS-Elements software (Nikon). A 7 x 2 μm z-stack was collected every 10 seconds for spindle pole separation, or every 20 seconds for fluorescence localization assays.

To assay chromosome dynamics, one-cell embryos expressing mCherry::H2b, GFP::H2b, or mSG::H2b were imaged on either the Nikon Ti2 (GFP::H2b and mSG::H2b), or with a Zeiss Axio Observer Z1 microscope equipped with a Yokogoawa CSU-X1 spinning disk and a QuantEM:512SC (Photometrics) EMCCD with a 60x 1.4 NA Plan Aprochromat Lens (Zeiss) using ZEN Blue microscopy software (Zeiss). High speed chromosome dynamics (chromosome span) were imaged with 6 x 2 μm z-stacks every 2 seconds.

Embryos (**Fig. 1B** and **Fig. 6C**) were also imaged on an Eclipse Ti2-E equipped with a CSU-W1 SoRA (Yokogawa) spinning disk head, a 60x 1.49 NA Plan Aprochromat Lens (Nikon), and dual Prime 95B CMOS cameras (Kinetix) at 12-bit settings using NIS-Elements software (Nikon). Images were acquired every 20 seconds using 7 x 2 μm z-stacks.

### Image analysis

ImageJ (FIJI) was used to process all microscope images. For spindle pole separation analysis, maximum intensity projections of GFP or mSG-tagged *tbg-1* and *his-11* were time-aligned to NEBD, which was defined as the frame when the diffuse nuclear histone signal equalized with the cytoplasm. Spindle length was then measured as the distance between the two centrosomes over time using ImageJ. Line scans of mNG-tagged *tpxl-1* in combination with mCherry-tagged *his-11* or *knl-1* were performed on maximum intensity projections of the 4 z-slices centered through the metaphase plate at −20s relative to anaphase onset. Anaphase onset was defined as the first time point in which chromosomes could be distinguished as two separate masses based on HIS-11 and TPXL-1 signals. A 50 pixel-wide line scan was drawn perpendicular to the long axis of the metaphase plate and centered on the chromosome midline. The fluorescence of the line was then measured by plotting the profile in ImageJ. Local background was then measured by extending the line by 2 μm and subtracted from the fluorescence intensity profile. Background subtracted values were then normalized to the peak intensity (the highest of the two peaks in KNL-1::mCherry) and plotted as a function of distance along the line scan.

Kinetochore and chromatin recruitment assays were performed on maximum intensity projections of the 4 z-slices fully encompassing the chromosome signal. A box was traced around chromosomes and kinetochores to measure chromatin and kinetochore signal respectively, and integrated density was recorded using Image J. Local background was calculated by extending the box by five pixels on all sides and subtracting the integrated density and area from the initial box. The integrated chromatin or kinetochore signal was then calculated by subtracting the background signal. This was repeated every 20s through anaphase onset for endogenously labeled SKA-1 dynamics, and 20s prior to anaphase onset for TPXL-1 and AIR-1 measurements.

Chromosome segregation analysis was performed on maximum intensity projections of mSG::H2b, mCherry::H2b, or GFP::H2b signals. Segregation errors were scored if the fluorescence histone signal was visible between separating sister chromatids for at least 20s after anaphase onset.

Chromosome span was measured in movies that were filmed with 2s intervals at 7 x 2 μm z-stacks for mSG::H2b and mCherry::H2b. Stacks were maximum projected, background subtracted, and converted to 8-bit in ImageJ. A minimum bounding box was then fit to the edge of the H2b fluorescence signal (the boundary where the pixel value is 0), and the width of the bounding box was measured at every 2 second interval during chromosome congression between NEBD and anaphase onset.

### Protein expression and purification

Codon-optimized gene blocks encoding the CLD (residues 205-257) of TPXL-1 (WT and V233>D;R226>A) were purchased from GenScript and cloned into the UC Berkeley Macrolab vector 2C-T (Addgene; #29706), which encodes an N-terminal TEV protease–cleavable His6-maltose binding protein (MBP) tag. Plasmids were transformed into Rosetta 2 (DE3) pLysS *E. coli* competent cells (EMD Millipore) and overnight cultures were used to inoculate 1-liter cultures of 2XYT media at 37°C with shaking at 180 RPM. At OD600 = 0.5, protein expression was induced with the addition of 0.2 mM isopropyl β-d-1-thiogalactopyranoside (IPTG), the cultures were shifted to 20°C, and incubated an additional 16 hours. Cells were harvested by centrifugation, resuspended in lysis buffer (20 mM HEPES-NaOH, pH 7.5, 300 mM NaCl, 10% glycerol, 5 mM imidazole, 5 mM β-mercaptoethanol), lysed by sonication (Branson Sonifier), and the cell lysate clarified by centrifugation. The supernatant was loaded onto a nickel affinity column (QIAGEN Ni-NTA Superflow) in lysis buffer, washed with 20 mM HEPES-NaOH, pH 7.5; 300 mM NaCl; 10% glycerol; 5 mM imidazole; 5 mM β-mercaptoethanol and eluted in 20 mM HEPES-NaOH, pH 7.5; 300 mM NaCl; 10% glycerol; 500 mM imidazole; 5 μM betamercaptoethanol. Fractions were pooled and diluted to 100 mM NaCl using 20 mM HEPES-NaOH, pH 7.5; 10% glycerol; 5 mM β-mercaptoethanol before loading onto an anion-exchange column (HiTrap Q HP, #17-1154-01; Cytiva). Bound proteins were eluted with a linear gradient from 100 mM to 1 M NaCl. Pooled fractions were then run on a size-exclusion column (Superdex 200 16/600; Cytiva) and pooled fractions concentrated, and stored at −80°C.

### Nucleosome binding assays

Nucleosome core particles (NCPs) were reconstituted following published protocols (<u>Luger et</u> <u>al, 1999</u>). Briefly, lyophilized *X. laevis* histones H2A, H2B, H3, and H4 were purchased from the Histone Source at Colorado State University (https://histonesource-colostate.nbsstore.net/). Histones were unfolded with incubation and shaking for 1h at RT in unfolding buffer (6 M guanidine hydrochloride, 20 mM Tris–HCl, pH 7.5, 5 mM DTT) and mixed in equimolar ratio for 1 mg/ml final concentration. Histones were refolded into an octamer by dialysis in refolding buffer (2 M NaCl, 10 mM Tris–HCl, pH 7.5, 1 mM EDTA, and 5 mM β-mercaptoethanol). After refolding, histones were concentrated via centrifugation and loaded onto a size-exclusion column (Superdex 200) equilibrated in refolding buffer. Fractions were analyzed by SDS–PAGE, and fractions containing pure histones were pooled. For nucleosome reconstitution, the Widom 601 DNA sequence was amplified by PCR, purified, and concentrated. DNA was added to the histone octamer in 1:1.2 molar ratio and dialyzed in 1.4 M KCl, 10 mM Tris–HCl, pH 7.5, 0.1 mM EDTA, 1 mM DTT for 1 hour at 4°C. Low-salt buffer (10 mM KCl, 10 mM Tris–HCl, pH 7.5, 0.1 mM EDTA, 1 mM DTT) was slowly pumped into the high-salt buffer for a couple of hours and then replaced with low-salt buffer and dialyzed overnight. Nucleosomes were concentrated with a centrifugal filter and injected onto a size-exclusion column (Superose 6) in 20 mM HEPES pH 7.5, 20 mM NaCl, 0.5 mM EDTA, 1 mM DTT. Fractions were analyzed on a 6% acrylamide TBE gel (prerun in 0.5X TBE for 150 V for 1 hour at 4°C) run for 1 hour at 150 V at 4°C. The gel was stained with SYBR Gold (S11494; Invitrogen; 1:10,000 dilution), imaged with a ChemiDoc system, and pure nucleosomes were pooled and concentrated with a centrifugal filter.

For the EMSA analysis, recombinant, purified His6-MBP-TPXL-1 CLD (WT or NB^mut^) and 25 nM reconstituted NCPs were incubated for 15 min at room temperature in binding buffer (20 mM Hepes, pH 7.5, 25 mM NaCl, 10% glycerol, 0.5 mM β-mercaptoethanol). After adding sucrose to 5% final concentration, samples were loaded onto a 6% TBE gel, (prerun at 150V for 1 hour at 4°C), and run for 80 min at 120V at 4°C in 0.2X TBE. The gel was then incubated in SYBR Gold for 10 min in the dark. Gel was imaged on a ChemiDoc Imaging system (12003153; ChemiDoc).

### AlphaFold3 Structure Predictions

Structure predictions of TPXL-1 and the nucleosome core particle or an H2A-H2B dimer was performed using Alphfold3 (*20*). PAE plots were generated using the PAEViewer web server (*33*).The *C. elegans* core histone sequences employed in the model are: HIS-3 (H2A), HIS-29 (H2B), HIS-2 (H3), HIS-67 (H4); note that there are multiple genes encoding 100% identical core histone sequences in the *C. elegans* genome (16 genes for H2A, 15 genes for H3, 16 genes for H4; for H2B, there are 5 genes encoding 100% identical sequence to HIS-3, and 6 genes encoding H2Bs with one additional amino acid, an alanine, in position 2. Structural analysis and depictions were carried out in PyMol (*34*).

### Statistical analyses

Statistical analysis was performed with Prism 10 software (GraphPad). Statistical significance of fluorescence intensity averages was determined by a two-sided Mann-Whitney test. Definitions for p-values are given in the corresponding figure legends. Error bars for pole-pole distance measurements represent standard deviations. Error bars for fluorescence intensities are 95% confidence intervals. Error bars for chromosome span analysis represent standard error of the mean. Each experiment represents (n) number of embryos from a minimum of five different hermaphrodites per condition.

## Acknowledgments

The authors thank Esther Zanin (Friedrich-Alexander University, Erlangen, Germany) for sharing the transgene-based TPXL-1 replacement system (plasmids and strains), Jessica Feldman (Stanford University, Palo Alto, USA) for sharing the *C. elegans*-optimized mStayGold sequence, and members of the Oegema and Desai labs for helpful discussions.

## Supplementary Materials

**Fig. S1.**
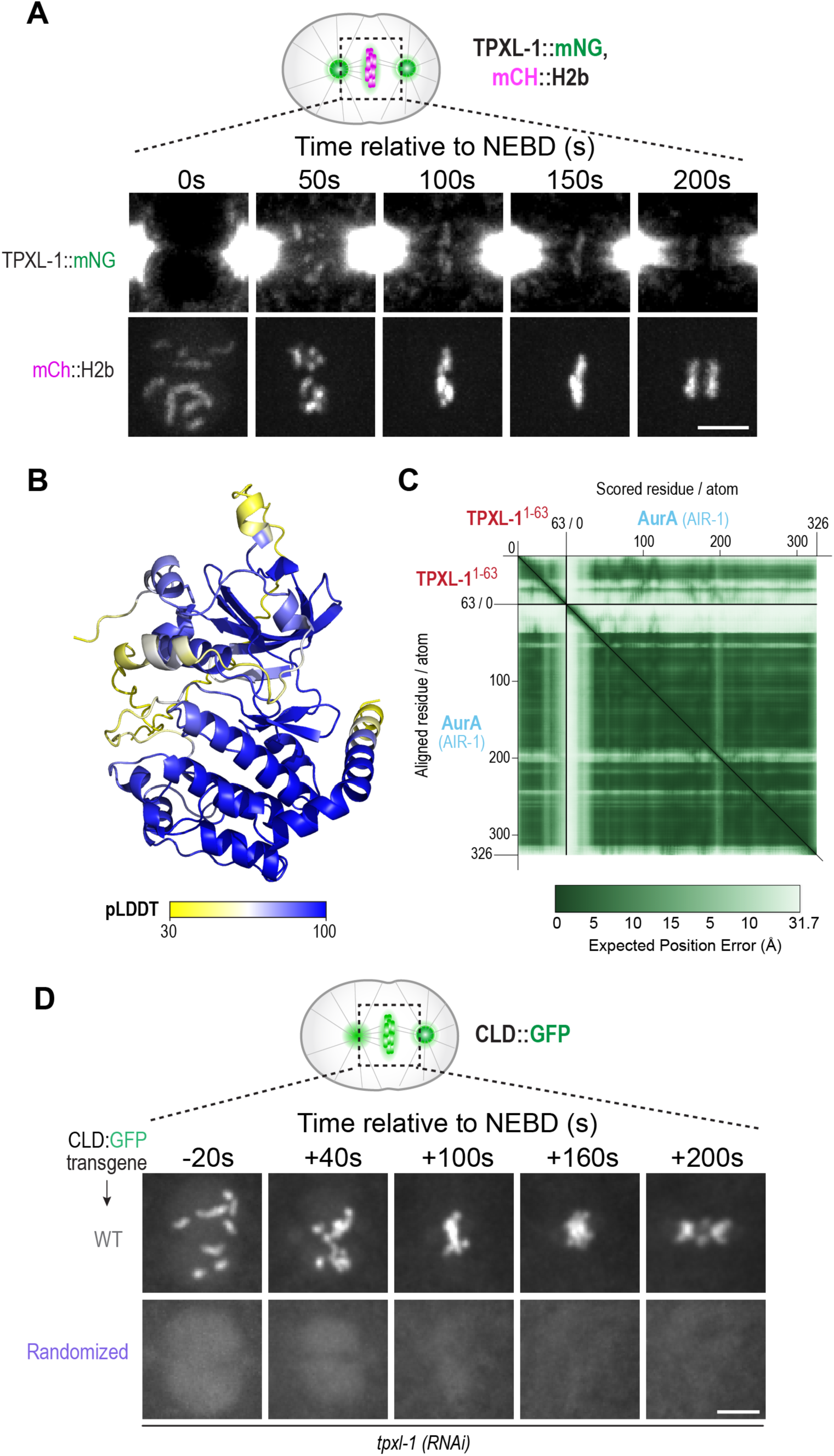
Alphafold model of TPXL-1–AurA interface and chromatin localization of TPXL-1::mNG over time. **(A)** Image panels from a timelapse movie of TPXL-1::mNG in an embryo co-expressing mCh::H2b. Contrast was enhanced in the TPXL-1::mNG channel to visualize the chromatin pool. Scale bar, 5 µm **(B)** AlphaFold3 predicted structure of TPXL-1^1-63^ and AurA (AIR-1), colored by confidence (pLDDT: predicted local distance difference test). **(C)** AlphaFold3 predicted aligned error (PAE) plot for TPXL-1^1-63^ and AurA (AIR-1) model, generated using the PAEViewer server.. **(D)** Image panels from timelapse movies of WT and Randomized TPXL-1 CLD, following endogenous TPXL-1 depletion. Scale bar, 5 µm. Image sequences in (*A*) and (*D*) were time-aligned relative to nuclear envelope breakdown (NEBD).

**Fig. S2.**
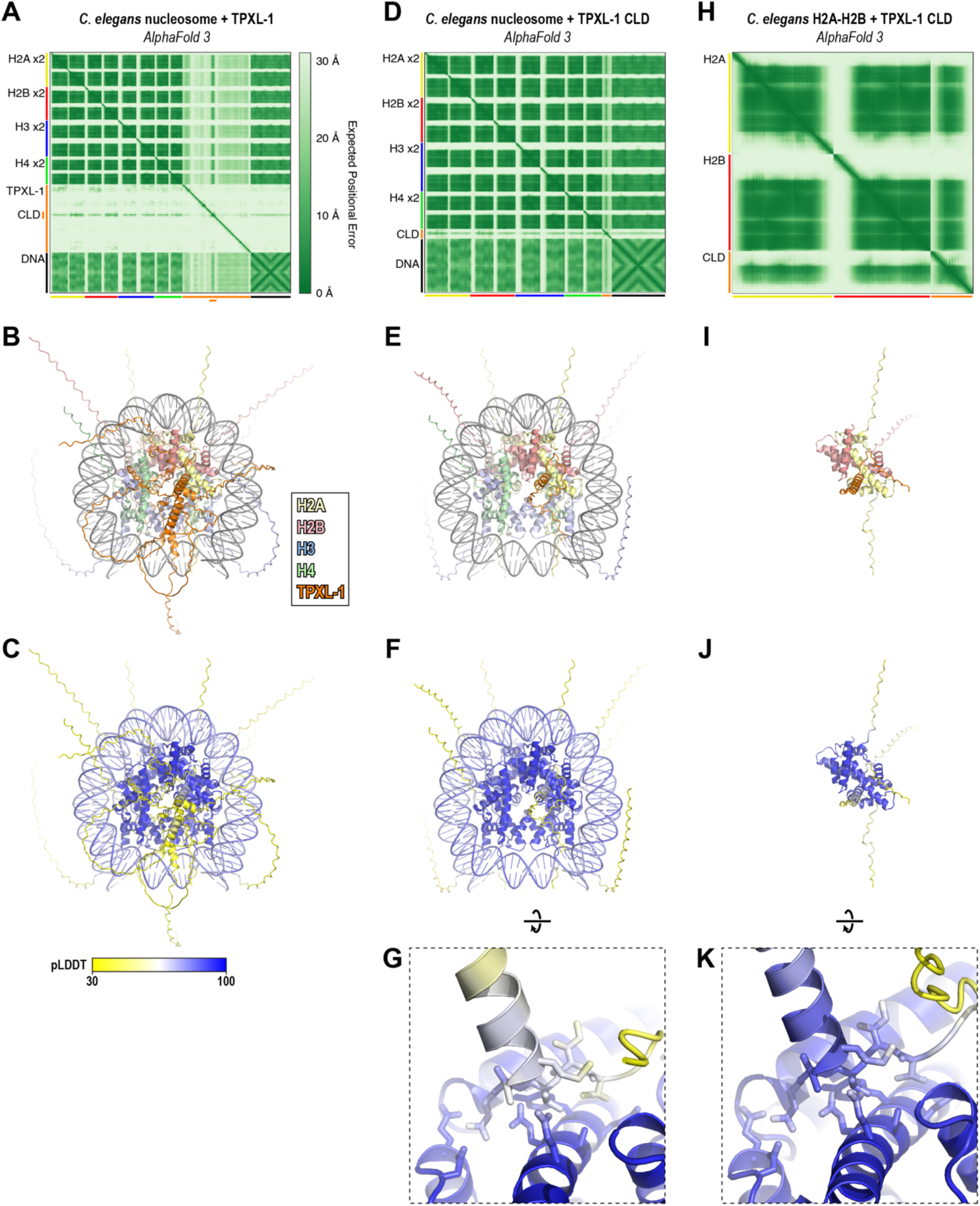
Alphafold3 models of TPXL-1 and the TPXL-1 CLD with the nucleosome core particle or an H2A-H2B dimer. **(A)** AlphaFold 3 predicted aligned error (PAE) plot for a prediction of a *C. elegans* nucleosome octamer (two copies each of histones H2A, H2B, H3, and H4) with full-length TPXL-1 and a double-stranded DNA (Widom 601 sequence). The *C. elegans* core histone sequences employed in the model are: H2A (HIS-3), H2B (HIS-29), H3 (HIS-2), H4 (HIS-67); note that there are multiple genes encoding 100% identical core histone sequences in the *C. elegans* genome. **(B)** AlphaFold 3 predicted structure of a *C. elegans* nucleosome octamer with full-length TPXL-1 and Widom 601 DNA. Histones are colored yellow (H2A), pink (H2B), blue (H3), and green (H4), and TPXL-1 is colored orange. **(C)** AlphaFold 3 predicted structure of a *C. elegans* nucleosome octamer with full-length TPXL-1 and Widom 601 DNA, colored by confidence (pLDDT: predicted local distance difference test). **(D)** AlphaFold 3 predicted aligned error (PAE) plot for a prediction of a *C. elegans* nucleosome octamer with the TPXL-1 CLD (aa 217-239) and a double-stranded DNA (Widom 601 sequence). **(E)** AlphaFold 3 predicted structure of a *C. elegans* nucleosome octamer with the TPXL-1 CLD and Widom 601 DNA, colored as in panel (*B*). **(F)** AlphaFold 3 predicted structure of a *C. elegans* nucleosome octamer with the TPXL-1 CLD and Widom 601 DNA, colored by confidence (pLDDT). **(G)** Closeup of the predicted interaction between TPXL-1 and the nucleosome acidic patch, from a prediction of a *C. elegans* nucleosome octamer with the TPXL-1 CLD and a double-stranded DNA (Widom 601 sequence), colored by confidence (pLDDT). View is equivalent to Fig. 3A (inset). **(H)** AlphaFold 3 predicted aligned error (PAE) plot for a prediction of a complex of *C. elegans* H2A and H2B with the TPXL-1 CLD. **(I)** AlphaFold 3 predicted structure of a complex of *C. elegans* H2A and H2B with the TPXL-1 CLD, colored as in panel (*B*). **(J)** AlphaFold 3 predicted structure of a complex of *C. elegans* H2A and H2B with the TPXL-1 CLD, colored by confidence (pLDDT). **(K)** Closeup of the predicted interaction between TPXL-1 and the nucleosome acidic patch, from a prediction of *C. elegans* H2A and H2B with the TPXL-1 CLD, colored by confidence (pLDDT).

**Fig. S3.**
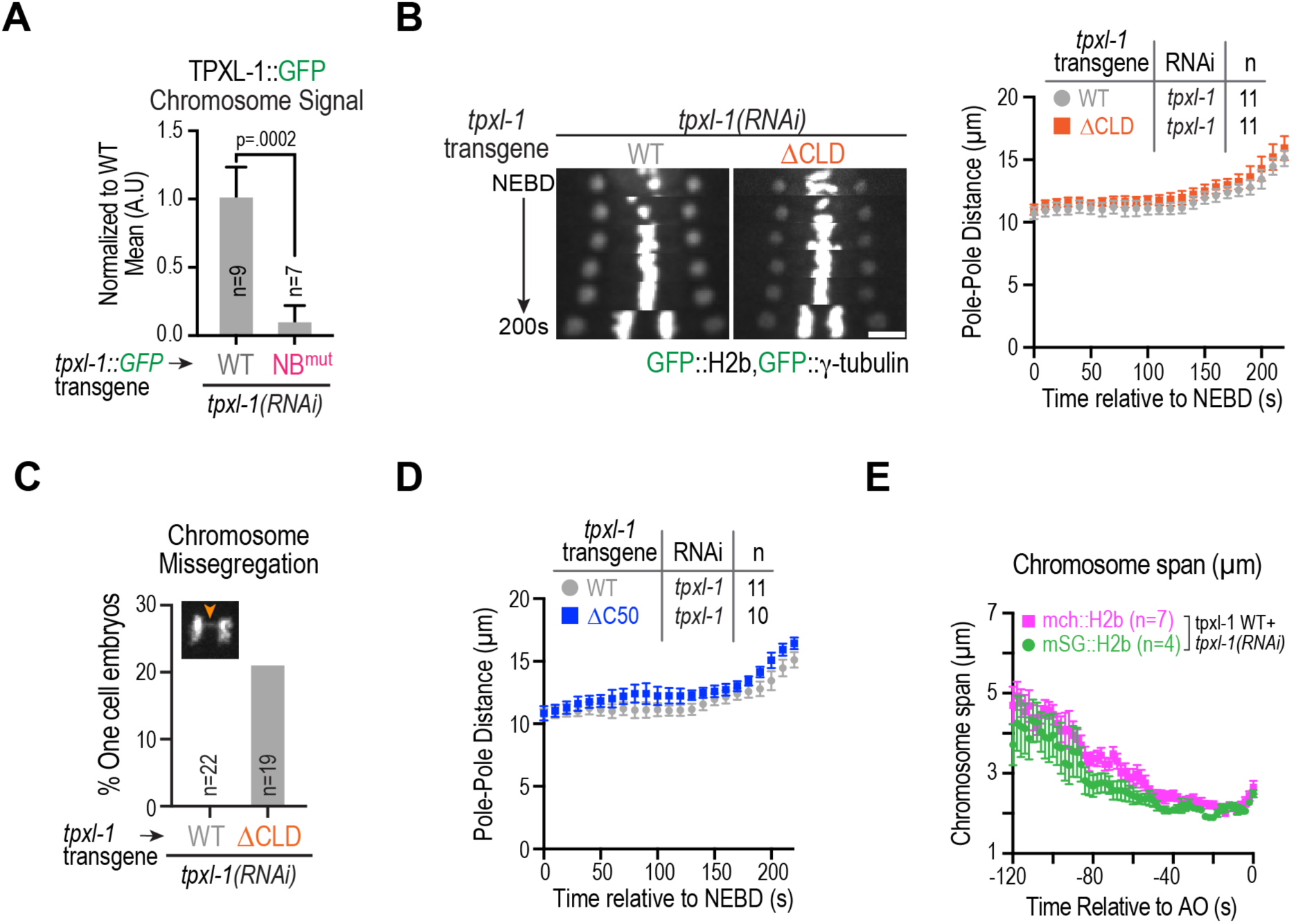
Additional localization and functional analysis of TPXL-1 variants. **(A)** Quantification of localization of NB^mut^ TPXL-1::GFP. *n* is the number of embryos imaged and analyzed. Error bars are the 95% CI. p-value (p=0.0002) is from a Mann-Whitney test. **(B)** (*left*) Kymograph of spindle region of embryos expressing GFP fusions to label chromosomes (H2b) and spindle poles (γ-tubulin). The indicated *tpxl-1* transgene insertions were present and endogenous TPXL-1 was depleted. Scale bar, 5 µm. (*right*) Graph of spindle pole separation over time for the indicated conditions. *n* is the number of one-cell embryos analyzed. Error bars are SD. **(C)** Graph plotting the percentage of one-cell embryos with chromatin bridges in anaphase for the indicated conditions. The embryos analyzed expressed GFP::H2b. *n* is the number of one-cell embryos analyzed. Note that the mCh::H2b and mSG::H2b analyzed for NB^mut^ TPXL-1 (Fig. 4C) are more sensitive at detecting anaphase chromosome bridges (mCh::H2b, because of significantly lower autofluorescence background; mSG::H2b because it is significantly more photostable and brighter than GFP::H2b, which is also prone to expression silencing when expressed from the transgene integrated on Chr V), which may account for the lower frequency of visible anaphase chromosome bridges scored for ΔCLD relative to NB^mut^ TPXL-1. **(D)** Graph of spindle pole separation over time for the indicated conditions. *n* is the number of one-cell embryos analyzed. Error bars are SD. WT data are the same as in (*B*). **(E)** Chromosome span analysis separating out mCh::h2b from mSG::H2b embryos in the TPXL-1 WT + *tpxl-1(RNAi)* condition. Error bars are SEM. These data were pooled and plotted in Fig. 5C: WT + *tpxl-1(RNAi)*.

**Fig. S4.**
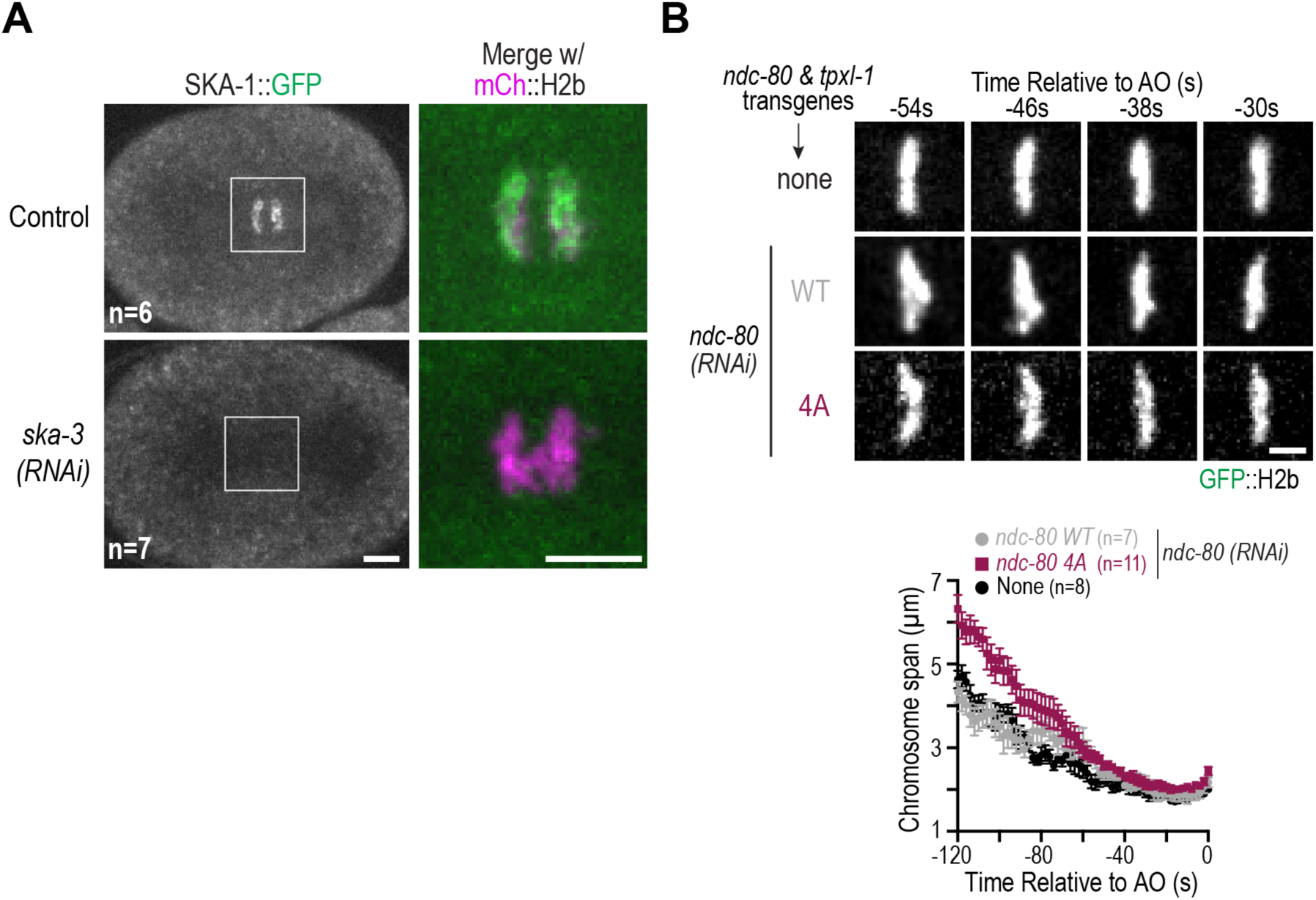
Confirmation of SKA complex removal and chromosome span analysis of the NDC-80 4A mutant. **(A)** Images of control and *ska-3(RNAi)* embryos with *in-situ* GFP-tagged SKA-1 that are expressing mCh::H2b. Chromosomal regions were boxed and magnified on the right. Scale bars, 5 µm for whole embryo and magnified panels. **(B)** Chromosome span analysis comparing WT to 4A NDC-80, following endogenous NDC-80 depletion. (*top*) Image panels from timelapse sequences for the indicated conditions. Scale bar, 2.5 µm. (*bottom*) Graph of chromosome span, analyzed as in Fig. 5C. Error bars are the SEM. Unlike WT NDC-80, NDC-80 4A mutant has chromosomes persisting near poles in early prometaphase which accounts for the initially larger chromosome span but they eventually congress to form a tight metaphase plate.

**Table S1.**
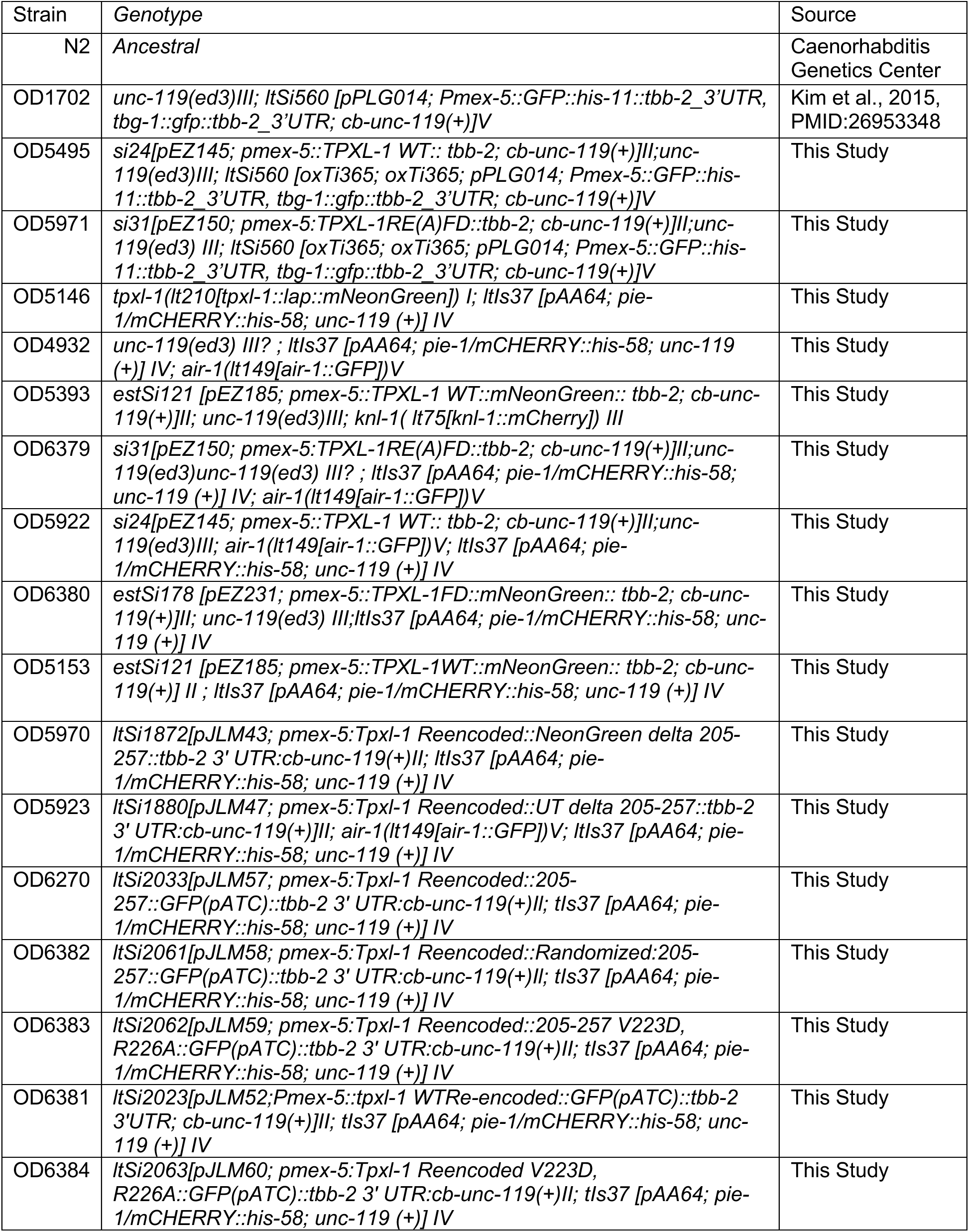
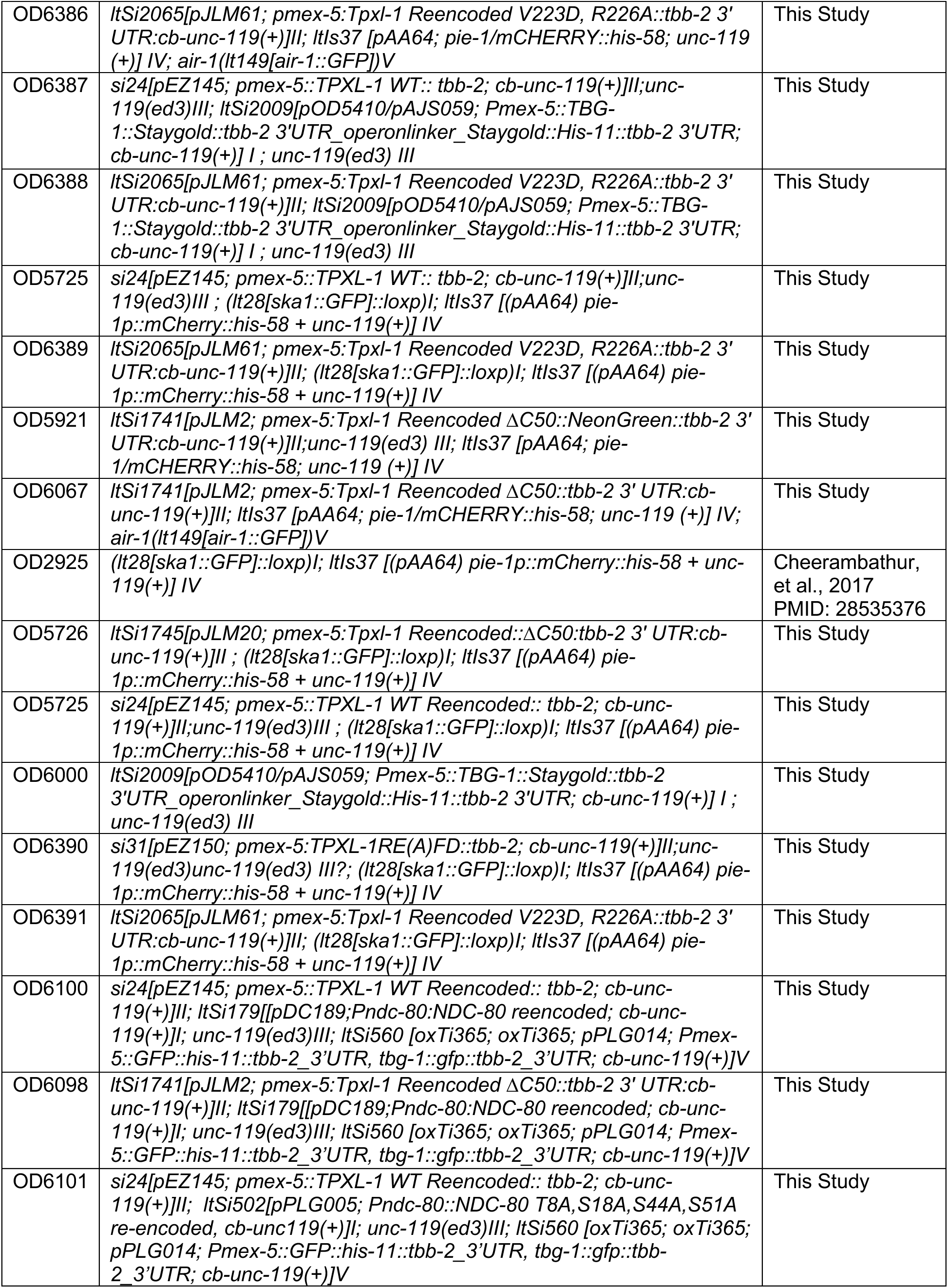
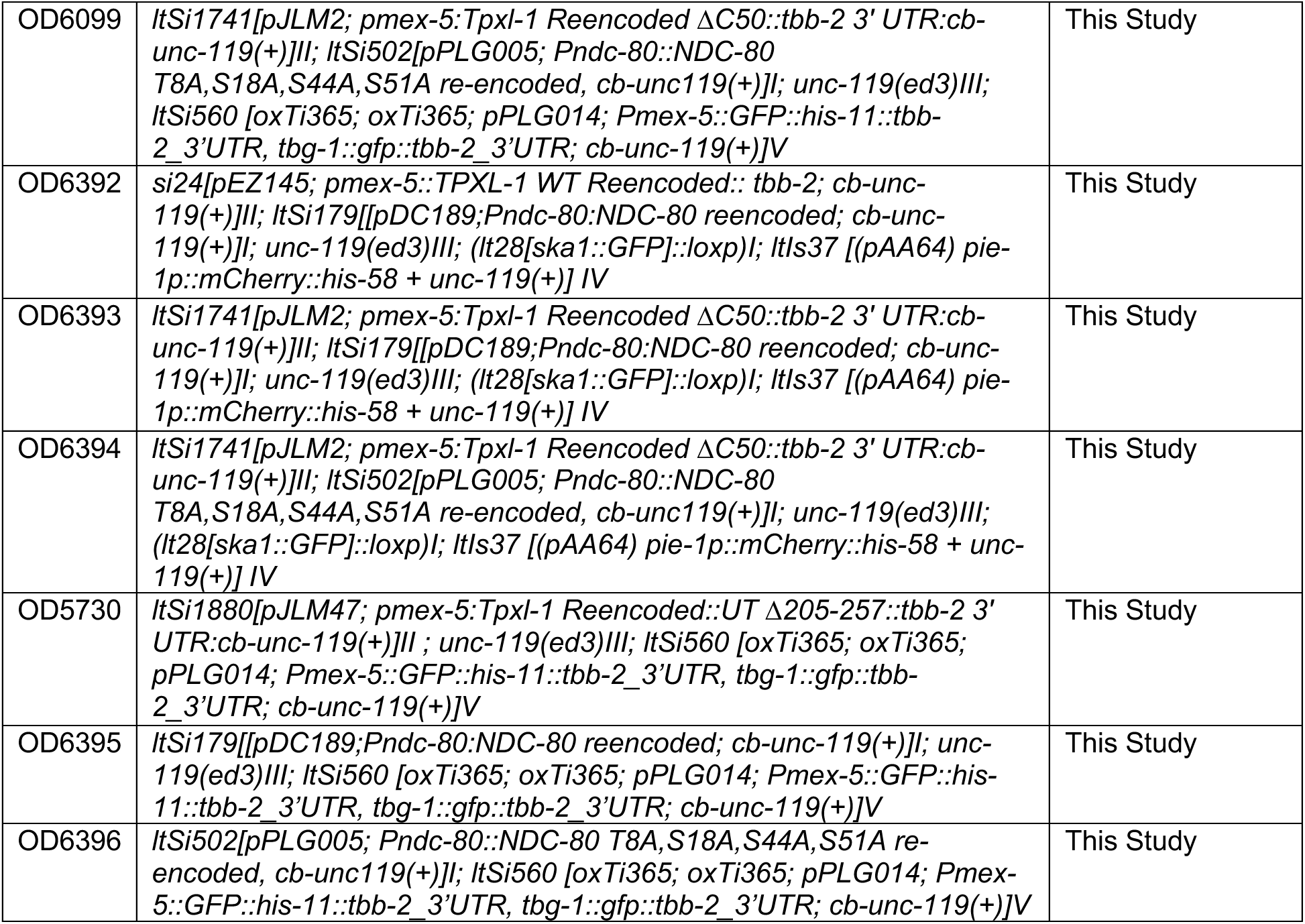
*C. elegans* strains.

**Table S2.**
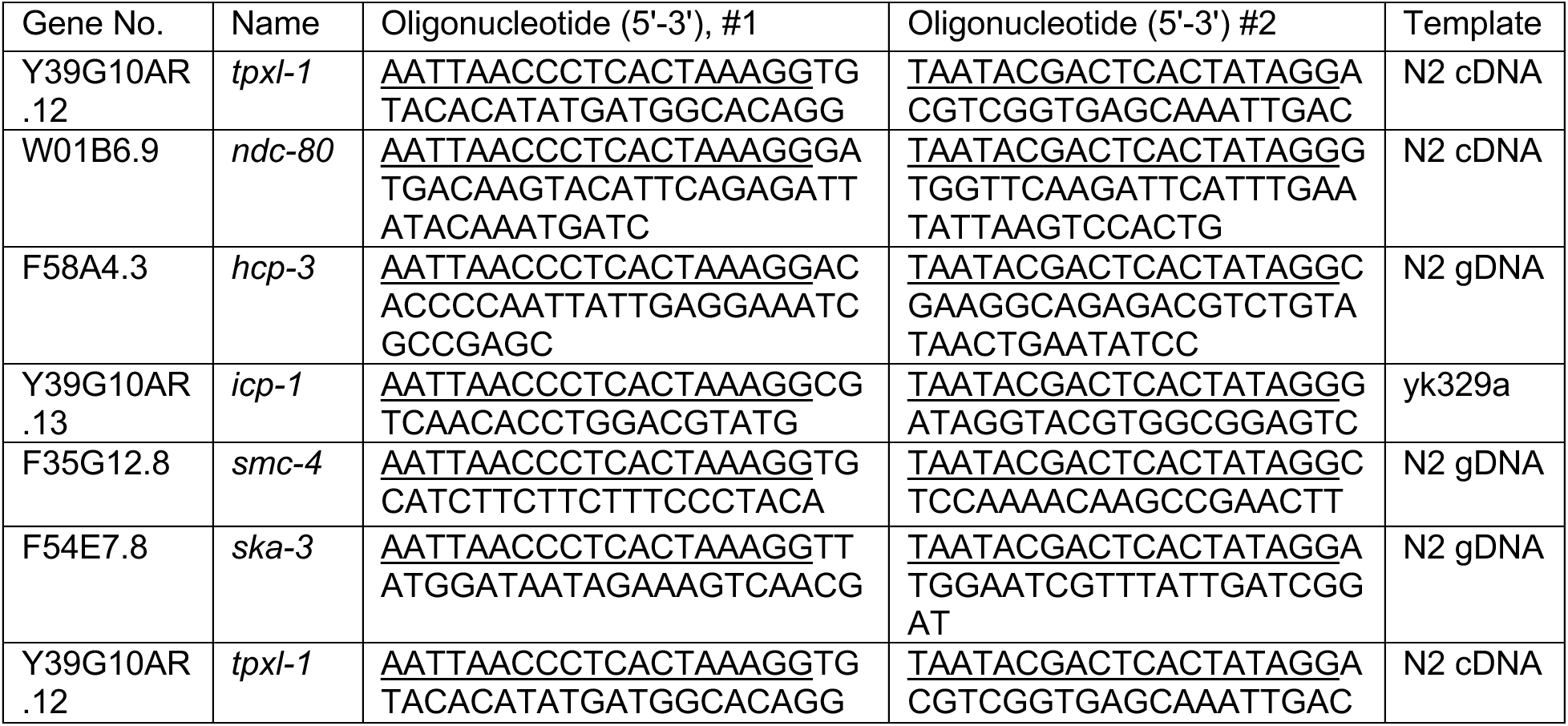
Primers for dsRNA synthesis.

